# Prediction of broad chemical toxicities using induced pluripotent stem cells and gene networks by transfer learning from embryonic stem cell data

**DOI:** 10.1101/2021.11.05.466718

**Authors:** Junko Yamane, Takumi Wada, Hironori Otsuki, Koji Inomata, Mutsumi Suzuki, Tomoka Hisaki, Shuichi Sekine, Hirokazu Kouzuki, Kenta Kobayashi, Hideko Sone, Jun K. Yamashita, Mitsujiro Osawa, Megumu K. Saito, Wataru Fujibuchi

## Abstract

The assessment of toxic chemicals using animals has limited applicability to humans. Moreover, from the perspective of animal protection, effective alternatives are also desired. Previously, we developed a method that combines developmental toxicity testing based on undifferentiated human embryonic stem (ES) cells (KhES-3) and gene networks. We showed that ≥ 95% accurate predictions could be achieved for neurotoxins, genotoxic carcinogens, and non-genotoxic carcinogens. Here, we expanded this method to predict broad toxicities and predicted the toxicity of 24 chemicals in six categories (neurotoxins, cardiotoxins, hepatotoxins, nephrotoxins [glomerular nephrotoxins/tubular nephrotoxins], and non-genotoxic carcinogens) and achieved high prediction accuracy (AUC = 0.90–1.00) in all categories. Moreover, to develop a testing system with fewer ethical issues, we screened for an induced pluripotent stem (iPS) cell line on the basis of cytotoxic sensitivity and used this line to predict toxicity in the six categories based on the gene networks of iPS cells using transfer learning from the ES cell gene networks. We successfully predicted toxicities in four toxin categories (neurotoxins, hepatotoxins, glomerular nephrotoxins, and non-genotoxic carcinogens) at high accuracy (AUC = 0.82–0.99). These results demonstrate that the prediction of chemical toxicity is possible even with iPS cells by transfer learning once a gene expression database has been developed from an ES cell line. This method holds promise for tailor-made safety evaluations using individual iPS cells.

## INTRODUCTION

To date, chemical toxicity studies have been primarily conducted by *in vitro* testing in cultured human cancer cell lines or in animals such as mouse, rat, and rabbit. However, because these methods differ from testing in "normal" human cells, their applications are limited^1^. In addition, the use of animals has become a major issue from the standpoint of animal welfare; in 2019, the U.S. Environmental Protection Agency announced that research studies using mammals as well as funding for mammal studies would be cut by 30% by 2025 and abolished by 2035^2^.

The embryonic stem cell test (EST) reported by Scholz et al. was the first to examine embryotoxicity *in vitro* using mouse fibroblasts, embryonic stem (ES) cells, and cardiomyocytes differentiated from ES cells; such developmental toxicity testing was previously performed only in animals^3,4^. Later, this method was approved as a scientifically valid alternative by the European Centre for the Validation of Alternative Methods (ECVAM) (https://tsar.jrc.ec.europa.eu/test-method/tm1999-01). However, the EST uses mouse cells, and species-specific differences must be clarified in order to use this approach to evaluate toxicity in human. Subsequently, another research group reported that in an EST based on a human cell system (hEST), changes in the expression of homologous neurodevelopmental genes were similar to those observed in the mouse system^5^.

In 2012, the U.S. Defense Advanced Research Projects Agency (DARPA) and the NIH invested a huge sum on a national project to promote the development of biomimetic systems, leading to rapid progress in the field^6^. These systems, which mimic the (adult) human body and are constructed by filling tissues created in individual compartments with culture fluid and connecting them together, are expected to be used in human toxicity testing systems as an alternative to animals. Currently, however, very little progress has been made in adapting these systems for application in developmental toxicity testing, and no evaluation method has been established for determining how accurately these systems mimic the function of the normal human body^7^. Realizing the practical application of these systems as high-throughput toxicity screening tools is likely to take several years.

For many years, the prediction of chemical toxicity has been carried out using a method based on the physicochemical parameters of the chemicals, referred to as the quantitative structure–activity relationship (QSAR)^8^. However, there is a limit to the predictive ability of QSAR. One reason is that the mechanism that actually induces toxic responses resides within the cell, so information about the chemical alone cannot predict such responses. In this regard, it should be possible to detect toxicity more accurately by obtaining information about the variation in gene expressions in cells, the so-called 'hardware' that mediates responses to chemicals. In addition, as stem cells differentiate, only the genes essential for that lineage is expressed and conserved through DNA methylation^9^, whereas in pluripotent stem cells, a very large number of genes are expressed, including transporters and transcription factors; consequently, pluripotent cells are superior to differentiated cells as a tool for comprehensively detecting toxic chemicals. In light of these considerations, we developed hEST-GN (human embryonic stem cell test with gene networks), a prediction method that uses information on feature gene networks based on massive gene expression datasets obtained by exposing human ES cells to toxic chemicals as input data for machine learning. Using this method, we achieved highly accurate predictions of developmental toxicity categories^10^.

In this study, we expanded the hEST-GN and found that the prediction of 24 toxic chemicals in broad toxicity categories, including adult toxicity, can be achieved with high accuracy. Furthermore, by selecting induced pluripotent stem (iPS) cells, which can be used as an alternative to ES cells in toxicity testing, and using a gene expression database created from ES cells, we were able to develop a method that can predict chemical toxicity using iPS cell data via transfer learning; this approach ameliorates the ethical issues related to ES cells. If this method could be further improved, it is likely that it could contribute to the development of tailor-made, individualized toxicity assessment/prevention using individual iPS cells, which would have enormous clinical value.

## RESULTS

### Development of an hEST-GN library for 24 chemicals and prediction using iPS

A schematic of the chemical assay is shown in **Fig. 1a**. The human ES cell line KhES-3 was exposed to a total of 24 chemicals in six toxicity categories [neurotoxins, hepatotoxins, cardiotoxins, two types of nephrotoxins (glomerular nephrotoxins and tubular nephrotoxins), and non-genotoxic carcinogens] at six concentrations, including vehicle (solvent) alone. The chemicals were carefully assigned to the toxicity categories by referring to previous reports (**Table 1**). Gene expression data were obtained by RNA-seq at two time points, 24 and 48 h, after exposure. At each time point, a principal component analysis (PCA) was performed using transcription factor genes, and a total of 20 genes from the top five PCs were extracted as feature genes. Using these genes, gene network libraries were created for each of the 24 chemicals using the Graphical Gaussian Model (GGM)^11^. Similarly, the screened iPS cells were subjected to RT-qPCR to obtain gene expression data for the same 20 genes and create gene network libraries. Next, using ES cell library labels, a chemical toxicity prediction system trained by both libraries from ES and iPS cells was developed via transfer learning^12^ using support vector machines (SVMs)^13^.

**Figure 1:**
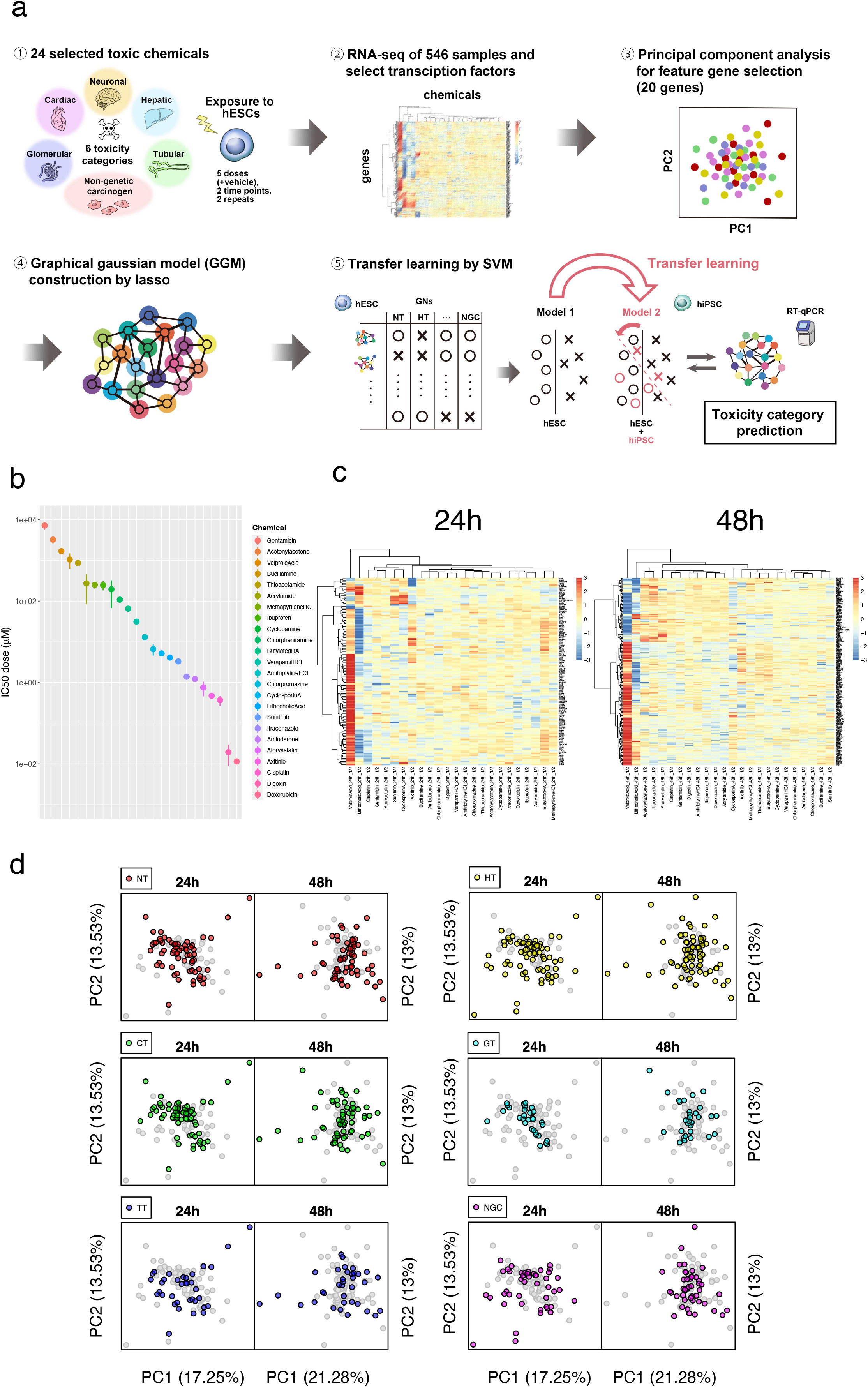
Construction of a gene expression database for 24 chemicals. (a) Schema of the chemical assay. hESC, human ES cells; hiPSC, human iPS cells. (b) IC50 for 24 chemicals. (c) Transcription factor genes differentially expressed following exposure to 24 chemicals at 1/2 dose. (d) PCA of 24 chemicals in six toxicity categories at two time points.

**Table 1.** List of 24 chemicals.

| Chemical name * | CAS RN | M.W. | NT | HT | CT | GT | TT | NGC |
| --- | --- | --- | --- | --- | --- | --- | --- | --- |
| Acetonylacetone | 110-13-4 | 114.14 | Nervous System Diseases [26] | NA | NA | NA | NA | NA |
| Acrylamide | 79-06-1 | 71.08 | Nervous System Diseases [26] | Liver Diseases [36] | NA | NA | NA | Group B2<br>https://www.epa.gov/iris |
| Amiodarone | 1951-25-3 | 645.3 | Cerebellar Diseases [27] | DILIrank: 8 (Most-DILI-Concern)<br>LiverTox: Likelihood score: A | Cardiotoxicity [41] | Glomerulonephritis, Membranous [54] | NA | NA |
| AmitriptylineHCl | 549-18-8 | 313.9 | Nervous System Diseases [28] | DILIrank: 5 (Less-DILI-Concern)<br>LiverTox: Likelihood score: A | Arrhythmias, Cardiac [42] | NA | NA | NA |
| Atorvastatin | 134523-00-5 | 558.6 | Hemorrhagic Stroke: [https://www.drugs.com/sfx/atorvastatin-side-effects.html] | DILIrank: 5 (Most-DILI-Concern)<br>LiverTox: Likelihood score: A | NA | NA | NA | NA |
| Axitinib | 319460-85-0 | 386.5 | NA | LiverTox: Likelihood score: E* | Cardiotoxicity [41]<br>Heart disease [43] | Proteinuria: [https://www.drugs.com/sfx/axitinib-side-effects.html] | NA | NA |
| Bucillamine | 65002-17-7 | 223.3 | NA | NA | NA | Nephrotic Syndrome [55] | NA | NA |
| ButylatedHA | 25013-16-5 | 180.24 | NA | NA | NA | NA | NA | Group 2B<br>Precancerous Conditions [66] |
| Chlorpheniramine | 132-22-9 | 274.79 | NA | DILIrank: 0 (No-DILI-Concern)<br>LiverTox: Likelihood score: E | NA | NA | NA | NA |
| Chlorpromazine | 50-53-3 | 318.9 | Parkinsonian Disorders [29] | DILIrank: 2 (Less-DILI-Concern)<br>LiverTox: Likelihood score: A | Cardiomyopathies [44] | NA | NA | NA |
| Cisplatin | 14913-33-8 | 300 | Peripheral Neuropathy: [https://www.drugs.com/sfx/cisplatin-side-effects.html] | DILIrank: 3 (Less-DILI-Concern)<br>LiverTox: Likelihood score: C | NA | NA | Acute Renal Failure [59] | Group 2A<br>http://ntp.niehs.nih.gov/ntp/roc/elevnth/profiles/s053cycl.pdf |
| Cyclopamine | 4449-51-8 | 411.6 | NA | NA | NA | NA | NA | NA |
| CyclosporinA | 59865-13-3 | 1202.6 | Abducens Nerve Diseases [30] | DILIrank: 7 (Most-DILI-Concern)<br>LiverTox: Likelihood score: C | Heart Diseases [45] | NA | Acute Kidney Injury [60] | Group 1<br>http://ntp.niehs.nih.gov/ntp/roc/elevnth/profiles/s053cycl.pdf |
| Digoxin | 20830-75-5 | 780.9 | Cognition Disorders [31] | DILIrank: 0 (No-DILI-Concern)<br>LiverTox: Likelihood score: E | Heart arrest [46] | NA | NA | Group 2B<br>https://monographs.iarc.who.int/list-of-classifications |
| Doxorubicin | 23214-92-8 | 543.5 | Spinal Cord Diseases [32] | DILIrank: 3 (Less-DILI-Concern)<br>LiverTox: Likelihood score: B | Arrhythmias, Cardiac [47] | Albuminuria [56] | NA | Group 2A<br>https://monographs.iarc.who.int/agents-classified-by-the-iarc/ |
| Gentamicin | 1403-66-3 | 477.6 | Hearing Loss [33] | LiverTox: Likelihood score: E | Heart Diseases [48] | NA | Kidney Tubular Necrosis, Acute [61] | NA |
| Ibuprofen | 15687-27-1 | 206.28 | Meningoencephalitis [34] | DILIrank: 3 (Less-DILI-Concern)<br>LiverTox: Likelihood score: A | Heart Septal Defects, Ventricular [49] | NA | Kidney Tubular Necrosis, Acute [62] | NA |
| Itraconazole | 84625-61-6 | 705.6 | NA | DILIrank: 8 (Most-DILI-Concern)<br>LiverTox: Likelihood score: B | Heart Failure [50] | NA | Kidney Diseases [63] | NA |
| LithocholicAcid | 434-13-9 | 376.6 | NA | Liver Cirrhosis [37] | NA | NA | NA | Precancerous Conditions [67] [68]<br>Cystic fibrosis [69] |
| MethapyrilineHCl | 135-23-9 | 297.8 | NA | Liver Neoplasms [38] | NA | NA | NA | Precancerous Conditions [70] |
| Sunitinib | 557795-19-4 | 398.5 | NA | DILIrank: 8 (Most-DILI-Concern)<br>LiverTox: Likelihood score: B | Arrhythmias, Cardiac [51] | Proteinuria [57] | NA | NA |
| Thioacetamide | 62-55-5 | 75.14 | NA | Hepatic Insufficiency [39]<br>Liver Cirrhosis [40] | NA | NA | NA | Group 2B<br>http://ntp.niehs.nih.gov/ntp/roc/elevnth/profiles/s172thio.pdf |
| ValproicAcid | 99-66-1 | 144.21 | Central Nervous System Diseases [35] | DILIrank: 8 (Most-DILI-Concern)<br>LiverTox: Likelihood score: A | Heart Septal Defects, Atrial [52] | NA | Kidney Diseases [64] | NA |
| VerapamilHCl | 152-11-4 | 491.1 | NA | DILIrank: 3 (Less-DILI-Concern)<br>LiverTox: Likelihood score: B | Ventricular Fibrillation [53] | Kidney Diseases [58] | Kidney Tubular Necrosis, Acute [65] | NA |

\* Chemical names are given by referring to PubChem (2021.4.20 ver.). Bold characters indicate positive toxicity.
NT, Neurotoxin; HT, Hepatotoxin; CT, Cardiotoxin; GT, Glomerular toxin (Renal toxin); TT, Tubular toxin (Renal toxin); NGC, Non-genotoxic carcinogen

### Gene expression response database of ES cells for 24 chemicals

To obtain the largest amount of data about the expression of genes that were perturbed by the 24 chemicals, ES cells need to be exposed to chemicals at the maximum concentration that does not cause an excessive degree of cell death. To this end, we first performed ATP assays and then plotted regression curves to calculate inhibitory concentrations (ICs) by carrying out serial dilutions of stock solutions containing the maximum soluble concentrations of the 24 chemicals. ICs in the range of 0.1% to 50%, at which cell death begins to be observed, were set as the maximum exposure concentration (**Table S1, Fig. S1**). The estimated IC50 and 95% confidence interval (CI) for each chemical are shown in **Fig. 1b** (**Table S2**). Serial dilutions were carried out, with the maximum exposure concentration set as 1/1 to obtain 1/2, 1/4, 1/8, and 1/16 dilutions, and a six-step exposure including vehicle alone was performed and repeated twice, yielding a total of 6 × 2 = 12 samples for each chemical. We collected RNA 24 and 48 h after exposure to the 24 chemicals, performed transcriptome analysis, and generated gene expression datasets for a total of 12 × 24 × 2 = 576 samples.

To examine the characteristics of the 24 chemicals at the level of differentially expressed genes (DEGs), we selected transcription factor–related genes (GO: 0006351) from the 576 datasets (4,032 genes). After log-normalization, batch effect elimination, and repeat merging, we generated DEG sets for which differences between each exposure data and vehicle values were significant (FDR < 0.01 and log_2_|FC| > 1) and presented them in a heatmap (**Fig. 1c, Fig. S2, S3**). According to this analysis, the number of DEGs was higher at 48 h than at 24 h for all concentrations, and over time, more genes were up- or down-regulated due to exposure to the chemicals. At both 24 and 48 h, valproic acid, a strong neurodevelopmental toxicant, elicited gene expression patterns that were clearly distinct from those of the other chemicals. Similarly, lithocholic acid, a mammalian bile acid and well-known carcinogen, yielded distinct expression patterns.

### Construction of gene networks of the 24 chemicals by the Graphical Gaussian Model (GGM)

To obtain feature genes used in the prediction, we performed PCA on the basis of the exposure data of each transcription factor gene, expressed as a log-fold–change (LFC) in expression relative to vehicle after log-normalization and batch effect elimination. There were 3,200 genes for the 24-h samples and 3,255 genes for the 48-h samples. For both exposure times, it was difficult to clearly separate the chemicals by the toxicity categories using two-dimensional PCA (**Fig. 1d**). Accordingly, we selected two genes with maximum positive and negative loading values, which were considered to contribute the most to the first to fifth PCs; at each time point, 20 genes were selected as feature genes (**Table S3**). Among the 20 selected genes in each group, only *ACTR3*^14^, which has been implicated in cell shape and motility, was common.

Using the 20 selected genes, we estimated sparse gene networks based on GGMs using an L1 graphical lasso for each chemical at each time point (i.e., 24 and 48 h) (**Fig. 2a, Fig. S4, Fig. S5**). The figures illustrate the estimated 190 partial correlation coefficients incorporated into the gene networks; edges with positive partial correlation values between two genes are shown in green, and edges with negative values are shown in red; the thickness, distance, and arrangement of the edges correspond to the degree of correlation between the two genes. Because these genes were obtained from the top five PCs that maximize the dispersion of the 24 chemicals using PCA, the estimated GGMs that describe the networks of all 20 genes differed considerably among chemicals, and it was difficult to classify the chemicals simply based on the network patterns as a whole. Therefore, for actual predictions, we decomposed the networks into their constituent edges rather than using them as a whole and used those with higher discriminative potential for training data as features. In other words, the partial correlation coefficients of the edges that are characteristic of the respective toxicity categories contributed to the SVM discrimination.

**Figure 2:**
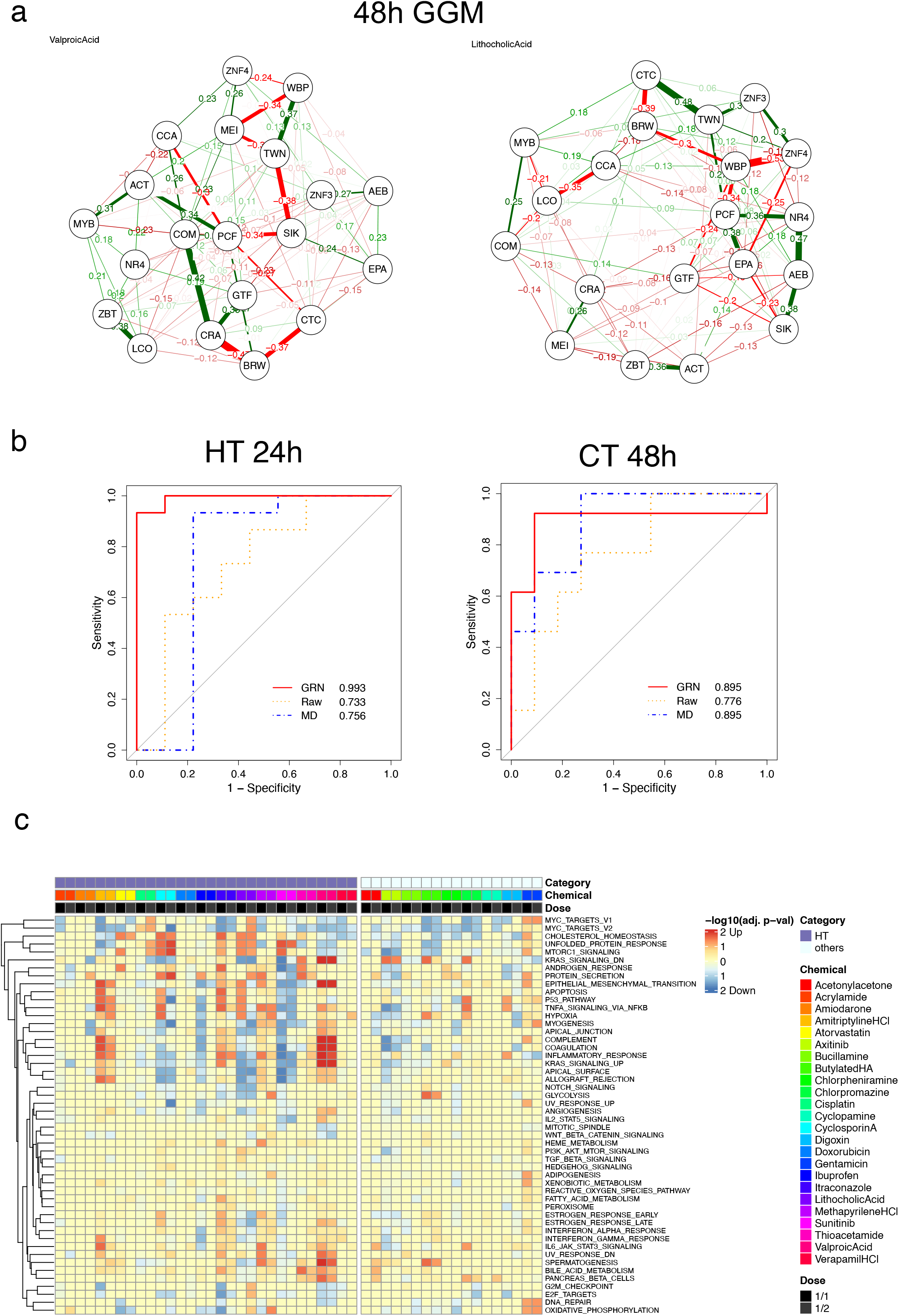
Prediction of six toxicity categories using KhES-3 cells. (a) Gene network representation of GGMs from KhES-3 cells. (b) ROC curves for the prediction of chemicals in two toxicity categories (c) Pathway analysis for hepatotoxins at 24 h and high-dose (1/1, 1/2 doses) samples.

### Prediction of six toxicity categories using KhES-3 cells and the GGM network

Using the 190 partial correlation coefficients in the GGM as input data, we predicted the toxicities of the 24 toxic chemicals in six categories using SVMs with leave-one-out-cross-validation (LOOCV). For the predictions, we followed the procedures described in a previous report on hEST-GN^10^ using four kernels (linear, polynomial, RBF, and maximum entropy) and increased the number of top features ranked by a t-test from 1 to 190. We also performed the prediction with the raw LFC values of the 3,200 (24 h) and 3,255 (48 h) transcription factor genes at each of the five concentrations relative to the vehicle-only expression. To compare the predictive accuracy, the same number of input data used for the GGM (i.e., up to 190 genes) was used as features. Predictions with the raw LFC values did not achieve a significantly higher prediction performance than the mean predictions using 10 uniform random numbers. On the other hand, in the prediction based on the GGM, the AUC values were ≥ 0.90 for chemicals in all toxicity categories, and because the prediction accuracy or AUC values were significantly high (p < 0.05), we concluded that prediction with high performance is possible. Predictions were performed separately at 24 and 48 h, but depending on the chemical, the time point at which a higher prediction accuracy could be obtained differed; thus, neither time point was considered particularly superior in terms of yielding a better prediction. Overall, these results demonstrated that hEST-GN based on ES cell gene networks allows for the prediction of not only developmental toxicity but also broad toxicity categories including adult toxicity.

In addition, to compare with predictions based on the QSAR theory, we generated 5,666 molecular descriptors including 3D descriptors (**Table S4**) and performed predictions using top 1 to 190 feature genes according to the aforementioned method. None of the six categories, except for tubular nephrotoxin (accuracy of 91.7%), gave a significantly high prediction result. The results of all predictions are presented together in a table and as ROC curves (**Table 2, Fig. 2b, Fig. S6**). These results suggest that chemical toxicity predictions that use the partial correlation coefficients of the GGM as features can achieve significantly higher accuracy than predictions based on gene expression values or QSAR. In the GGM-based prediction, the prediction accuracy for each of the 24 chemicals was examined from the SVM results. This analysis revealed that with respect to 16 chemicals (acetonylacetone, acrylamide, amitriptyline HCl, atorvastatin, bucillamine, chlorpheniramine, chlorpromazine, digoxin, doxorubicin, gentamicin, itraconazole, lithocholic acid, methapyrilene HCl, sunitinib, thioacetamide, and valproic acid), the prediction accuracy was 100% for all six categories at 24 and 48 h. On the other hand, for axitinib, cisplatin, and cyclosporin A, the prediction accuracy was 66.6%, suggesting that the prediction of these chemicals is difficult (**Table S5**).

**Table 2.**
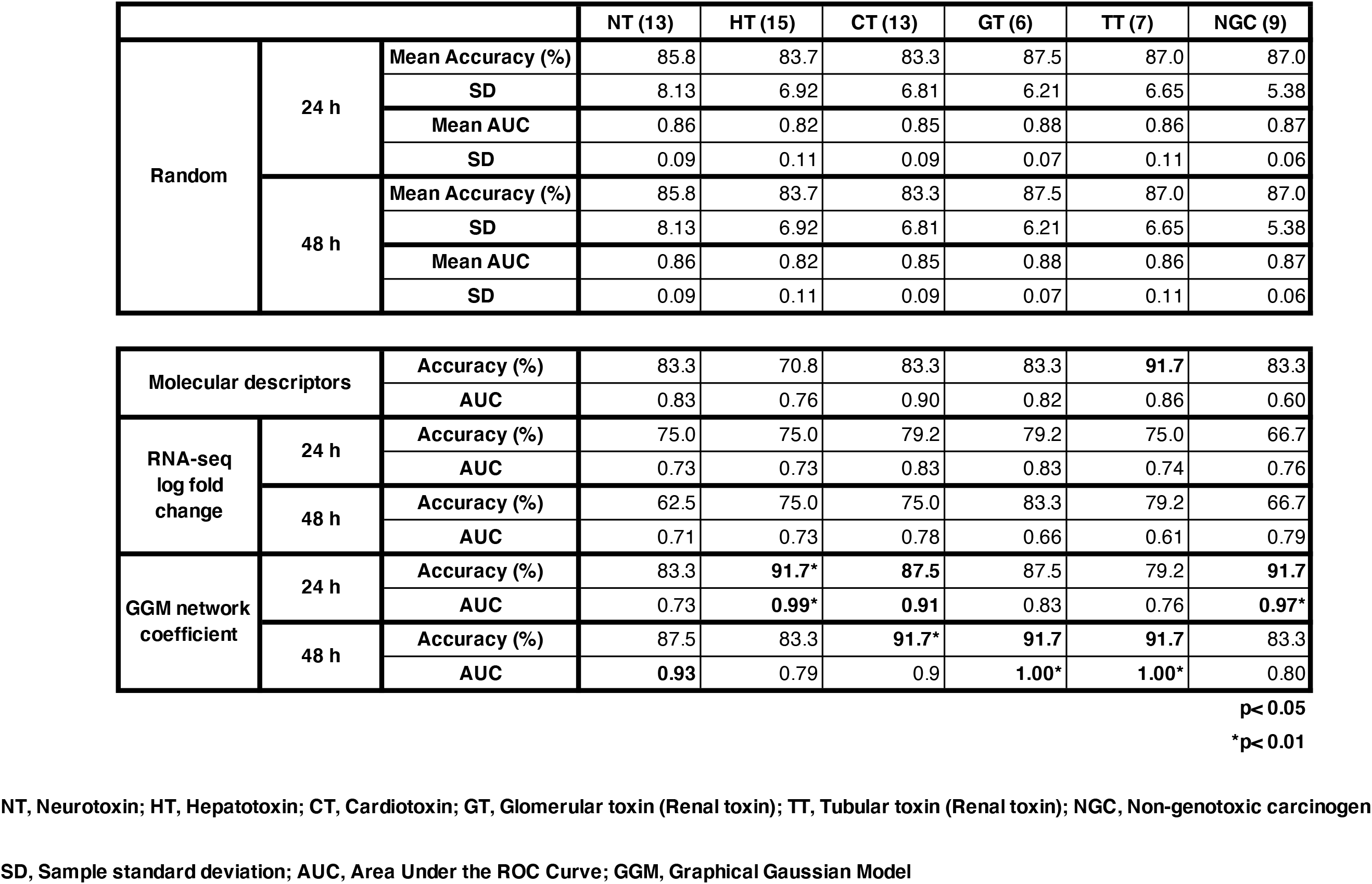
Summary of prediction performance for KhES-3.

### Pathway analysis of KhES-3 genes following exposure to 24 chemicals

To determine the effects of exposure to chemicals on biological pathways, we performed a Hallmark pathway analysis by Gene Set Enrichment Analysis (GSEA) using all genes. When performing this analysis, we divided the samples into high-dose (1/1 and 1/2 doses) and low-dose (1/8 and 1/16 doses). For hepatotoxins, cardiotoxins, and globular nephrotoxins, differences were observed in the types of pathways that were induced or suppressed in comparison with other toxic chemicals in the high-dose samples at 24 h (**Fig. 2c, Fig. S7**). In addition, the responses of ES cell genes to the toxic chemicals were diverse and dependent on the type of chemical to which they were exposed and not limited to specific pathways such as apoptosis. This observation suggests that it is possible to predict toxicity categories on the basis of perturbed pathways that can be detected by transcription factors. On the other hand, differences in concentrations or among categories that may have been present at 48 h were not as pronounced as those at 24 h. However, analyses using available pathways based on human knowledge accumulated in the past provide limited information. Instead, the computational extraction of feature genes from the PCA of all genes without bias and predictions based on their GGM networks are likely to be more effective.

### Selection of iPS cells as an alternative to human ES cells

The results of the present and previous studies suggest that hEST-GN can predict not only developmental toxicity but also broad toxicity categories with high performance. However, there are still hurdles to overcome, including ethical issues, before this system can be generally and widely accepted as a toxicity test. Accordingly, to make iPS cells a possible alternative to hEST-GN, we performed pre-screening by comparing ATP assays with ES cells exposed to 20 toxic chemicals across a wide range of categories. As candidates, we used the top 20 cell lines selected from among Japanese male cell lines^15^ derived from healthy individuals, which had been examined and ranked in terms of their differentiation potential into the three germ layers. For exposure concentration, we adopted the IC50 that was determined using the KhES-3 cell line and examined the toxicity response of human iPS cells. Among the candidate cell lines, we selected the top three cell lines with well-correlated growth rates at IC50 (HPS4138, HPS4234, and HPS4046) and confirmed the correlation coefficients of the growth rates at IC50 with KhES-3 using 20 of the 24 toxic chemicals investigated in this study. HPS4138 had the highest value of 0.94; accordingly, this cell line was used for the predictions as an alternative to ES cells (**Table S6**).

### Prediction of chemicals in six toxicity categories using HPS4138 iPS cells

For HPS4138, which was selected by screening, we performed ATP assays with the 24 chemicals (**Fig. S8, Table S7, Table S8**). As in the case of ES cells, the cells were exposed to vehicle alone or five concentrations obtained by serial dilutions of stock solutions containing the maximum exposure concentration (i.e., the maximum value between IC0.1 and IC50 that did not cause an excessive degree of cell death); this experiment was repeated twice. Gene expression data for 20 genes selected from KhES-3 cells at 24 and 48 h were obtained by qRT-PCR, and GGMs were created for each of the 24 chemicals based on LFC values relative to the vehicle. Partial correlation coefficients were used for the prediction, as in the case of ES cells. In the prediction, data were created by integrating iPS cell data with ES cell data as well as by transductive transfer learning, in which toxicity category labels in the ES cell data were used for the learning to allow for category prediction using iPS cells. Assessment was performed by LOOCV, similarly to the predictions made using ES cells only. Chemicals in all categories except cardiotoxins and tubular nephrotoxins yielded AUC values from 0.82 to 0.99, and the accuracy or AUC was significantly higher than for results obtained with uniform random numbers. Thus, although this approach was not perfect, the results of the prediction using HPS4138 were very accurate for most toxicity categories (**Table 3**). The summary of toxicity category predictions for the 24 chemicals using HPS4138 are shown in **Fig. 3**. Prediction was difficult for butylated HA, but satisfactory for the other chemicals (**Table S9**). These results demonstrate that if a gene expression database for toxicity responses could be created with ES cells, chemical toxicity prediction using iPS cells would also be possible by means of transductive transfer learning. Our findings also raise the possibility of achieving practical applications of toxicity testing using standardized or individualized iPS cells in the future. A description of transductive transfer learning is available on our web site at https://hipst-gn.stemcellinformatics.org.

**Figure 3:**
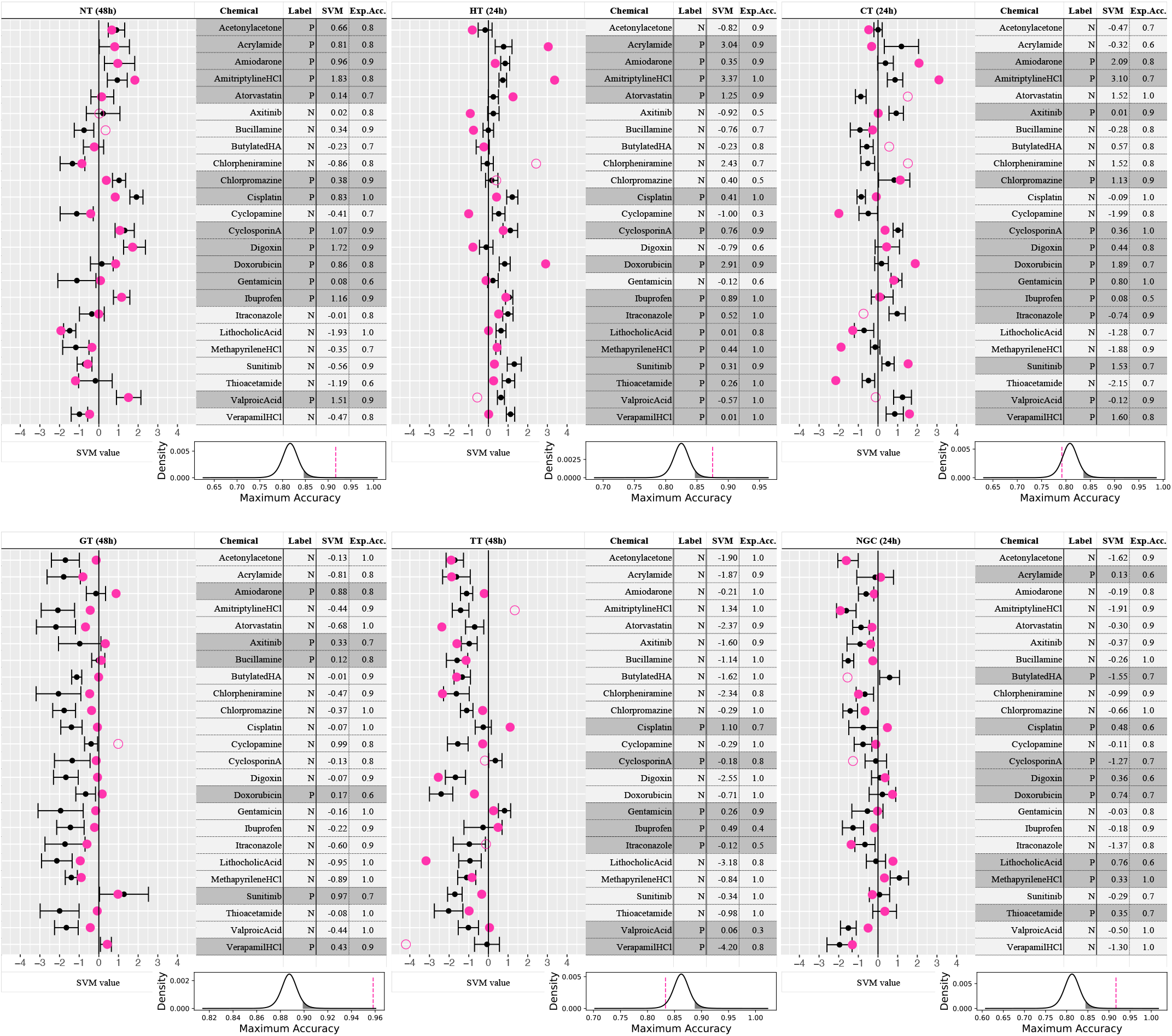
Summary of toxicity category prediction for 24 chemicals using HPS4138 cells. Blue dots indicate predicted SVM values of iPS cell data, and filled and open markers indicate true and false predictions, respectively. Black dots and bars indicate the means ± S.E.M. of SVM values for random data. In the tables, the label columns contain prior knowledge regarding whether the chemical shows toxicity (P: positive) or not (N: negative). The SVM column contains the SVM values of the iPS cell data. The expected accuracy (Exp.Acc.) columns contain the prediction accuracy using random data. The graphs below the tables show the probability distribution of the prediction accuracy using random data. Black lines indicate the probability density estimated using the t distribution (degrees of freedom = 9). Black shaded areas represent the upper 5%. Blue dashed lines indicate the prediction accuracy using iPS cell data.

**Table 3.**
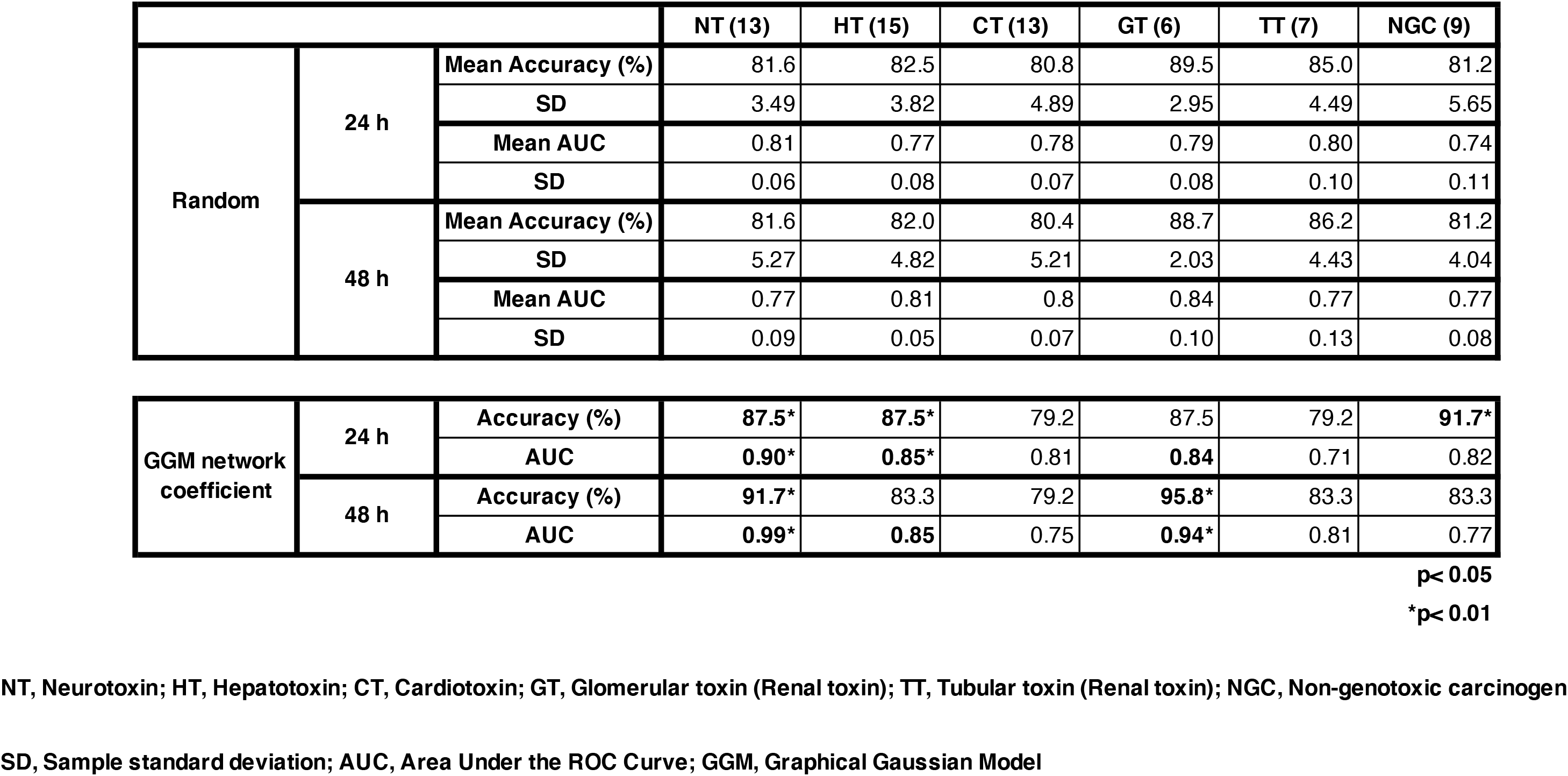
Prediction for HPS4138 cells.

## DISCUSSION

Here we presented a proof-of-concept study that enables a toxicity hazard assessment using human iPS cells and transfer learning based on the transcription factor gene network libraries made from the gene expression data of human ES cells exposed to 24 chemicals for 6 categories.

In this paper, we clarified that i) human ES cells are sufficient to detect not only developmental toxicities during embryogenesis but also broad toxicity categories such as adult toxins (neurotoxin, cardiotoxin, hepatotoxin, and glomerular and tubular nephrotoxins) and non-genotoxic carcinogens; ii) the chemical toxicity prediction using transcription factor gene networks of human ES cells shows an AUC = 0.90–1.00, which is significantly more accurate than predictions based on the QSAR theory or from raw gene expression data; iii) there exist differences in the biological pathways affected by the toxicity categories, suggesting the mechanisms that underlie the transcription factor networks that control the pathways may be used to predict toxicity categories; and iv) the gene network data from properly screened human iPS cells can successfully, although not perfectly, predict the toxicity categories at significant accuracies once the toxicities are learned by transfer learning using the models based on human ES cell data only.

Various alternative methods using pseudo-human systems, such as differentiated human cell lines (HepG2, MCF-7, HeLa, etc.), have been reported^16^ since animal protection has become a higher priority in research. However, these lines are often derived from cancer or immortalized cells and thus have limited use^17^. On the other hand, primary cells, which are assumed to resemble natural states in the human body, show batch-to-batch variability^17^ and are difficult to collect at sufficient amounts. Furthermore, it is difficult to extrapolate toxicity tests on some cell types to other target cell types due to differences in cytotoxicity tolerance^18^. Performing a multi-target toxicity prediction system based on stem cells, as we propose here, provides a more valid prediction of toxicity to a larger range of cell types.

Toxicological assessment using the transcriptome is frequently used by the U.S. EPA, Tox21 project and in Europe. Particularly, New Approach Methodologies (NAMs), which are any technology, methodology, approach or combination thereof that can be used to provide information on chemical hazards and risk assessments that avoids the use of intact animals^19^, are often directional concepts using transcriptomics with other omics or traditional toxicology methods. Our study indicated that transcription factor gene networks exist in a master layer of biological pathways to activate molecular initiation events (MIEs), in which the initial chemical trigger starts an adverse outcome pathway (AOP) via DNA-binding, receptor activation, or a disturbance of cellular / organelle systems^20^, thus revealing toxicity reactions. Recent studies have endorsed the idea that transcription factors of signal receptors might play a role in interfacing outside signals, such as aryl hydrocarbon or androgen, to activate toxicological AOPs in HepaRG^TM^ cells^21^. In our system, we used 20 transcription factor genes from 5 PCs due to limited resources and cost, but it should be possible to customize the set of genes and PCs to reflect more accurately the specific AOPs in the endpoint organ. To pursue a full coverage of endpoint organs and AOPs, RNA-seq analysis and a library of all 4,033 transcription factor genes for the test chemicals at low cost are needed.

Previous systems using QSAR theory depend on the information of chemicals only and are thus inapplicable to mixtures such as food, Chinese or herbal medicines, and other compounds to assess toxicity as a whole. Our hEST / hiPST-GN system detects the cellular toxic events of these mixtures, providing the prediction of holistic cellular reactions, making it a resource for industries that mix independent chemical ingredients, including cosmetic, air-conditioner, and automotive companies who need to assess the toxicity of mixtures in their final or intermediate products, media, emissions, detergents, etc. In fact, more than 100 members from a wide variety of industry-government-academia fields are involved in our non-profit consortium (scChemRISC). By developing products from candidate substances that are predicted to have little toxicity, our system will contribute to industry not only for efficiency but also for human health.

Recently, toxicity reaction differences due to ethnicity, or genome haplotypes, have been widely reported due to the globalization of foods and products among countries. For example, catechin, which is contained in green tea and is widely consumed in Asian countries, is reported to induce severe liver injury in the United States^22^. The CiRA Foundation (Kyoto, Japan) has announced myiPS cells, a project in which individuals can have their own iPS cells generated and banked. Our hiPST-GN system could allow a tailor-made chemical toxicity assessment for these cells to detect individual differences in toxic tolerance for different substances. Ideally, it also has the potential to reduce medical accidents if myiPS cells could be used to diagnosis whether a medication is toxic before receiving the treatment^17,23^. In general, by performing a battery of toxicity assessments using multiple iPS cell lines from individuals with various haplotypes, our system may contribute to reducing toxicity accidents often caused by a small number of test samples of limited genomic variances.

In conclusion, the largest advantage of our hEST / hiPST-GN is the ability to perform toxicity hazard assessments for multiple endpoints with high accuracy in a short amount of time and a low cost. We believe that our system will greatly benefit research that will be affected by the lost funding for mammal studies designated to happen in 2035 by the U.S. EPA.

## EXPERIMENTAL PROCEDURES

### Cell culture experiments

The KhES-3 cell line was established at and provided by Kyoto University^24^. The protocol of this study was reviewed by the Ethics Committee of CiRA in accordance with the "Guidelines for Derivation and Utilization of Human Embryonic Stem Cells" by the Ministry of Education, Culture, Sports, Science and Technology, Japan. The iPS cell lines were established from healthy Japanese donors at CiRA, Kyoto University, and were approved for use by the Ethics Committee of Kyoto University.

Since it has been reported that the toxicity of antioxidants such as catechin is suppressed in the presence of albumin^25^, maintenance culture was carried out for all cell lines including human ES cells using albumin-free Essential 8 Medium (Thermo Fisher Scientific) in six-well feeder-free culture dishes coated with 5 μg/mL vitronectin (VTN-N; Thermo Fisher Scientific). When seeding the cells, 10 μM CultureSure^®^ Y-27632 (FUJIFILM WAKO) was added, and medium exchange on day 1 and thereafter was performed without Y-27632.

### Selection of toxic chemicals

Twenty-four chemicals were selected and mainly included neurotoxins, hepatotoxins, cardiotoxins, nephrotoxins, and non-genotoxic carcinogens (**Table 1**). The presence or absence of toxicity was determined mainly on the basis of information regarding toxicity and assorted disorders and diseases available at PubChem (https://pubchem.ncbi.nlm.nih.gov/). With respect to neurotoxicity, among the chemicals previously reported in the literature, those having only developmental toxicity were classified as ‘negative,’ as the present study targeted adult toxicity. In addition, when determining the presence or absence of hepatotoxicity, chemicals with DILI rank ≥ 3 were considered hepatotoxic chemicals; with regard to others, those with reliable reports of liver diseases were considered hepatotoxic. As for cardiotoxicity, chemicals that have been reported to be associated with heart disease were considered cardiotoxic. With regard to the kidney, due to its diverse and complex structure, the area of damage was divided into two sites, namely, the glomeruli and renal tubules. Chlorpheniramine and cyclopamine were the chemicals judged to be completely ‘negative’ and belonged to none of the toxicity categories examined in the present study. On the other hand, 19 chemicals had multiple overlapping toxicities, whereas acetonylacetone, bucillamine, and butylated HA had only one toxicity^26-70^.

### Chemical exposures and determination of IC logistic model equations

DMSO or water was used as solvent (vehicle) for the 24 chemicals based on known information (https://pubchem.ncbi.nlm.nih.gov/). For each of the 24 toxic chemicals, a stock solution was prepared with the highest soluble concentration. First, in order to perform the ATP assay to determine the exposure doses for testing, we performed 10 serial three-fold dilutions of the stock solution; the prepared exposure solution was added to the cells (i.e., exposure) so that the concentration in the medium was 0.1% of the exposure solution. The cells were cultured on 96-well black/clear flat bottom TC-treated plates (Falcon), 8,000 cells were seeded, and the medium was exchanged on day 1. The cells were exposed to chemicals on day 2. No medium exchange was performed after exposure, and the ATP assay was performed 48 h after exposure. For 100 μL of culture medium, 100 μL of CellTiter-Glo^®^ Luminescent Cell Viability Assay (Promega Corporation) was added, and emitted light was measured with a 2104 EnVision Multilabel Plate Reader (PerkinElmer). From four luminescence measurements for 10 concentrations and a blank as described above, regression analysis was performed by fitting the three-parameter log-logistic model with the R-4.0.5 drc package, and IC0.1 and IC50 values were obtained (**Tables S1 and S2**). IC50 values were plotted using the ggplot2 package **(Fig. 1b)**.

### RNA-seq analysis

On day 2 after seeding, the cells were exposed to the chemicals. For each of the 24 chemicals, the exposure dose was set between IC0.1 and IC50 depending on the degree of cell death. Using this value as the maximum exposure dose, five serial two-fold dilutions (1/1, 1/2, 1/4, 1/8, 1/16) were performed, and a total of six doses including a solvent-only control (vehicle) were used. After exposure, no medium exchange was performed, and samples were obtained at two time points (24 h and 48 h) with two repeats, i.e., a total of 24 × 6 × 2 × 2=576 samples. After RNA purification using an RNeasy Mini Kit (QIAGEN), sequence libraries were prepared for each sample using TruSeq Stranded mRNA Library Prep/TruSeq RNA Single Indexes Set A & Set B (Illumina, Inc.). For sequencing, high-throughput sequencing was performed using HiSeq4000 (Illumina, Inc.). We used bowtie-2.2.5 with the option “--very-sensitive-local” to map the obtained Illumina reads to Ensemble GRCh38r100 human cDNA and ncRNA sequences, added up the reads for each gene using MAPQ ≥ 1 transcript, and obtained average counts of 28,652,809 and 28,291,682 reads for each sample at 24 h and 48 h, respectively. From these, we selected only transcription factor-related genes included in the Gene Ontology GO:0006351 using BioMart (4,032 genes). These genes were filtered using the statistical analysis language R-4.0.5 package edgeR^71^ (https://www.r-project.org/) with the filterByExpr function min.count = 30, min.total.count = 0, and then normalized to log2 counts per million (logCPM) using the voom function. Furthermore, the removeBatchEffect function was used to eliminate batch effects.

### Differentially expressed gene analysis and principal component analysis

At 24 h and 48 h, for a total of 122 groups including 120 conditions (five concentrations each for 24 chemicals) and two solvent conditions (DMSO or water), we used a linear model fitting with the lmFit function of the limma package^72^ in R and moderated t-statistics with eBayes to analyze DEGs with respect to gene expression levels in terms of the LFC between 24 chemicals and their corresponding solvent^73^, and created a heatmap of genes with an LFC > 1 and FDR (false discovery rate) < 0.01 using the pheatmap package in R (**Fig. 1c, Fig. S2)**. In addition, using the LFC values obtained for 120 conditions, PCA was performed for each of the two time points (24 h, 48 h) using the prcomp function in R (**Fig. 1d, Fig. S3**).

### Gene network construction by Graphical Gaussian Model (GGM)

Based on results obtained in the aforementioned PCA, a total of 20 genes (two genes each with the top positive and negative loading values in the first to fifth PCs) were used to construct gene networks. To estimate the GGM for each of the 24 chemicals, we used the aforementioned LFC values to calculate the sparse partial correlation coefficient network with L1 graphical lasso using EBICglasso in the R package qgraph (https://cran.r-project.org/web/packages/qgraph/qgraph.pdf)^74^. For model fitting, regular BIC with gamma = 0 was used, and regularization of sparsity was tried 1000 different ways with nlambda = 1000 for estimation. In the estimation, in order to avoid the problem of covariance matrices failing to be positive definite, calculations were performed with checkPD=FALSE (**Fig. 2a, Fig. S4, Fig. S5**).

### Prediction by support vector machine (SVM)

The SVM program and protocol used in the present study were adopted according to the report of Takahashi et al.^75^ Four kernel functions were used: linear, polynomial, RBF, and maximum entropy, and parameter types and combinations were calculated according to the above report^75^. In the calculation, 190 values of partial correlation coefficients among 20 genes in the GGM for each of the 24 chemicals described above were used as input data, and using LOOCV, the genes were ranked using the two-sample t-test (two-sided) in each iteration, and the maximum accuracy and the maximum AUC to achieve the maximum accuracy were recorded, varying the number of values from 1 to 190. To statistically evaluate the maximum accuracy and AUC, 24 × 190 uniform random numbers were generated 10 times, and the maximum accuracy with the maximum AUC were recorded in a similar fashion. One-sample t-test (one-sided) was performed with the average values and standard deviations of maximum GGM accuracies and AUCs obtained from these 10 attempts. For comparison, LFC data for the transcription factor genes at five concentrations before calculating the GGM (24 × 5 × 3,200 or 3,255 values in total) were used as input data, and the maximum accuracy and the corresponding AUC value were recorded in a similar manner. In addition, to compare with predictions based on QSAR, 5,666 molecular descriptors were created using alvaDesc (Affinity Science) (https://www.alvascience.com/alvadesc/). We obtained information from PubChem DB (https://pubchem.ncbi.nlm.nih.gov/) 3D Conformer data regarding 20 of the 24 chemicals; for the four other chemicals for which information could not be obtained from PubChem DB (cisplatin, cyclosporin A, digoxin, and gentamicin), the SMILES format was converted into 3D molecular descriptors using CORINA Classic (https://www.mn-am.com/online_demos/corina_demo) and entered into alvaDesc.

### Gene set enrichment analysis

We created LFC data for all genes by performing the same preprocess as for the transcription factor genes (21,650 and 22,298 genes for 24 h and 48 h, respectively) and divided them into high-dose (1/1/, 1/2) and low-dose (1/8, 1/16) groups to perform GSEA for each group using the R package fgsea^76^. As for the gene sets used, among the MSigDB Collections provided by GSEA (https://www.gsea-msigdb.org/gsea/index.jsp), we used 50 hallmark gene sets. The heatmap was generated with FDR-adjusted p values obtained using the fgsea package.

### Selection of HPS4138 iPS cells

The measurement of the differentiation potential to the three germ layers was performed according to a previous report^15^, where we examined the expression ratio of two marker genes for each layer (*PAX6*, *SOX2*, *BRA*, *NCAM*, *SOX17*, and *FOXA2*) by means of fluorescence activated cell sorting. Among the ranked Japanese male cell lines derived from healthy individuals, we used the top 20 cell lines according to their total ratios^15^. These cell lines were kept in maintenance culture with StemFit AK02N medium (Ajinomoto) and then cultured for two passages in maintenance culture using Essential 8 Medium (Thermo Fisher Scientific) as in the case of KhES-3. Among the 20 cell lines, one underwent cell death, and the remaining 19 were subjected to pre-screening using 20 chemicals with a wide range of toxicities (valproic acid, cyclopamine, acrylamide, acetonylacetone, chlorpromazine, chlorpheniramine, atorvastatin, amiodarone, verapamil HCl, dimethoate, arsenic trioxide, quinidine, axitinib, doxorubicin, gentamicin, ibuprofen, lithocholic acid, thioacetamide, butylated HA, and methapyrilene HCl) by comparing the ATP assay results with those for ES cells. The details of the ATP assay are described above. For exposure concentrations, the IC50 determined with KhES-3 was used, and the growth rate of human iPS cells was examined. Among the candidate cell lines, the top three cell lines (HPS4138, HPS4234, and HPS4046) whose growth rates at IC50 correlated well with that of KhES-3 were selected, and again, the growth rate at IC50 was confirmed by the ATP assay using 20 of the 24 toxic chemicals examined in the present study (**Table S6**) to select the cell line with the largest correlation coefficient, i.e., HPS4138.

### Gene expression data from HPS4138 by RT-qPCR

The five serial exposure concentrations of the 24 chemicals for iPS cells were determined by the ATP assay, as described above with respect to ES cells. For each of the 20 genes at 24 h and 48 h used for the construction of the GGM for ES cells, the primer sequence pair was designed using Primer 3 (version 0.4.0) based on the human cDNA sequence data obtained from Ensembl GRCh38r100, and the obtained primer sequence pair was synthesized (Hokkaido System Science). We confirmed whether the target PCR product could be obtained from the primer sequence pair according to the product size determined by electrophoresis. On day 2 after seeding, HPS4138 cells were exposed to the 24 chemicals. With regard to exposure concentrations, time points, and repeat experiments, we followed the experiments performed with ES cells. After purification using an RNeasy Mini Kit, RNA was transcribed to cDNA using a PrimeScript^TM^ RT Reagent Kit (Perfect Real Time) (TaKaRa), and the synthesized primers for qRT-PCR and KAPA SYBR Fast qPCR Kit (KAPA BIOSYSTEMS) were used to perform qRT-PCR with StepOnePlus (Applied Biosystems). ΔC_T_ values of the resulting genes were obtained by subtracting from the C_T_ values the value of the internal control (*GAPDH* gene). The average value of two repeated measurements at each concentration was determined. In addition, for solvents (DMSO and water), ΔC_T_ values were obtained by subtracting the value of *GAPDH* gene, and the average of all values was determined. The difference, ΔΔC_T_, between the ΔC_T_ value at each concentration and that of the solvent was determined and is referred to as LFC. Covariance matrices among the 20 genes were calculated and used as input data in the construction of the GGM.

### Transductive transfer learning by SVM

A total of 380 edges in the GGM of ES cells and the GGM of iPS cells were used to perform the prediction of chemicals in the six toxicity categories under conditions similar to the SVM protocol described above. For learning, labels of the 24 chemicals in ES cells were provided, whereas none of the labels of the 24 chemicals in iPS cells were provided (i.e., zero); prediction was performed via transductive transfer learning. As in the case of ES cells, prediction was also performed by replacing with uniform random numbers 10 times only the values of the 24 x 190 input data for iPS cells, and similar to ES cells, the one-sample t-test (one-sided) was used to assess the maximum accuracy and the corresponding AUC value.

## Supporting information

Exteded Data Figures S1-S8

## DATA AVAILABILITY

The source code of analysis program and the data supporting the findings of this study are available from the corresponding author upon reasonable request. The RNA-seq data reported in this paper have been deposited in NCBI’s Gene Expression Omnibus and are accessible through GEO series accession number GSE188203.

## SUPPLEMENTAL INFORMATION

Supplemental Information includes eight extended data figures and nine supplemental tables and can be found with this article online at:

## AUTHOR CONTRIBUTIONS

WF conceptualized the research and led the project. JY drafted the manuscript. JY and KK provided the stem cell data. JY, TW, WF, and JKY analyzed the data. HO, KI, MS, TH, SS, HK, and HS selected the toxic chemicals. MO, and MKS provided the iPS cell lines.

## ACKNOWLEDGEMENTS

The authors deeply appreciate Prof. Shinya Yamanaka for kindly advising on the selection of iPS cell lines. The authors also deeply appreciate Dr. Peter Karagiannis for kindly reviewing the manuscript. This work was partially supported by Kyowa Kirin Co., Ltd., Shiseido Co. Ltd. (for only iPS cell line screening), and the Core Center for iPS Cell Research, Research Center Network for Realization of Regenerative Medicine (20bm0104001h0008/21bm0104001h0009) and the Program for Intractable Diseases Research utilizing Disease-specific iPS cells (19bm0804001), Japan Agency for Medical Research and Development (AMED).

