## Supplementary material for "Prediction of broad chemical toxicities using induced pluripotent stem cells and gene networks by transfer learning from embryonic stem cell data": Exteded Data Figures S1-S8

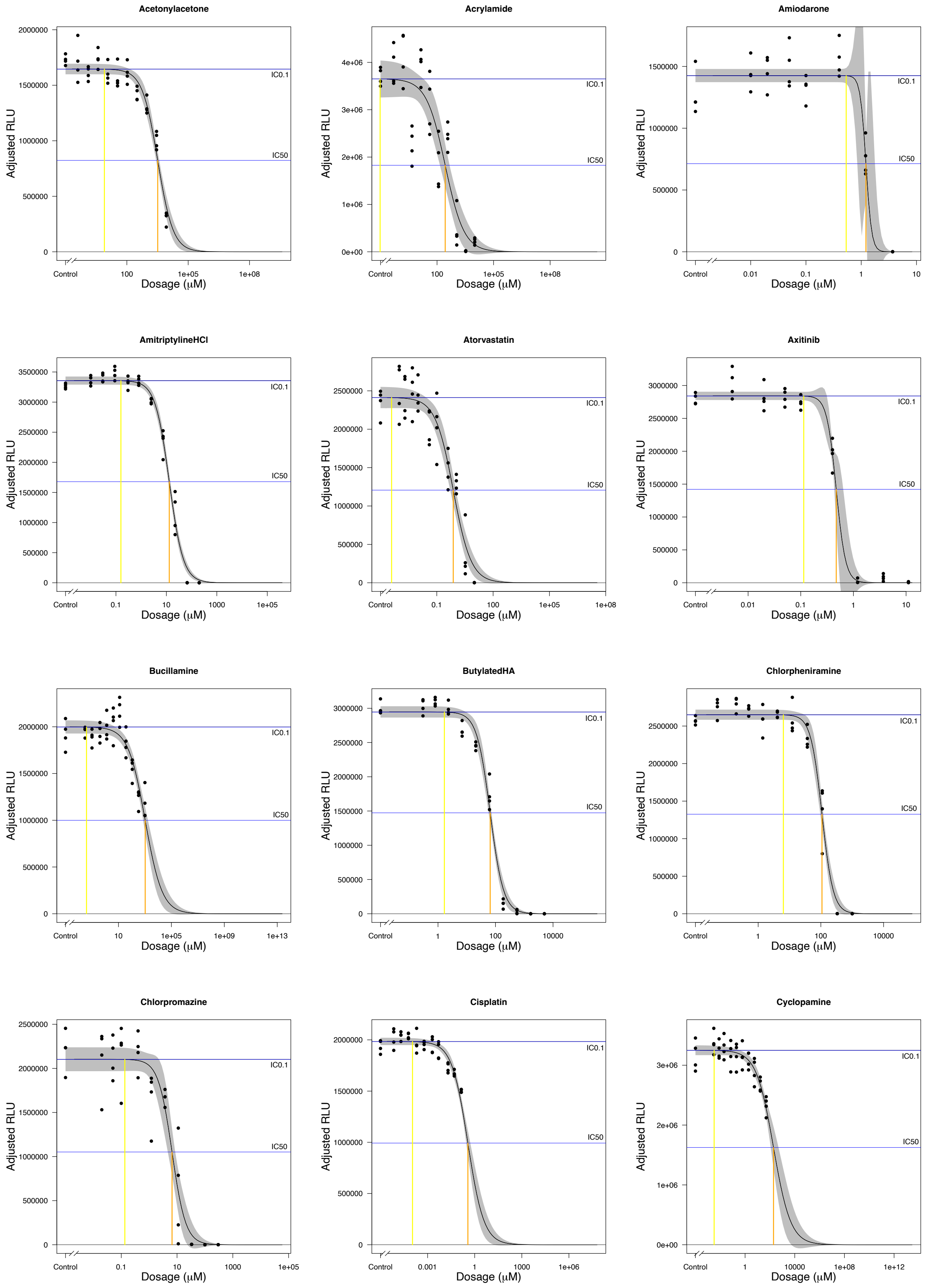
Acetonylacetone
Acrylamide
Amiodarone
AmitriptylineHCl
Atorvastatin
Axitinib
Bucillamine
ButylatedHA
Chlorpheniramine
Chlorpromazine
Cisplatin
Cyclopamine
Adjusted RLU
Dosage (μM)
Control
IC0.1
IC50

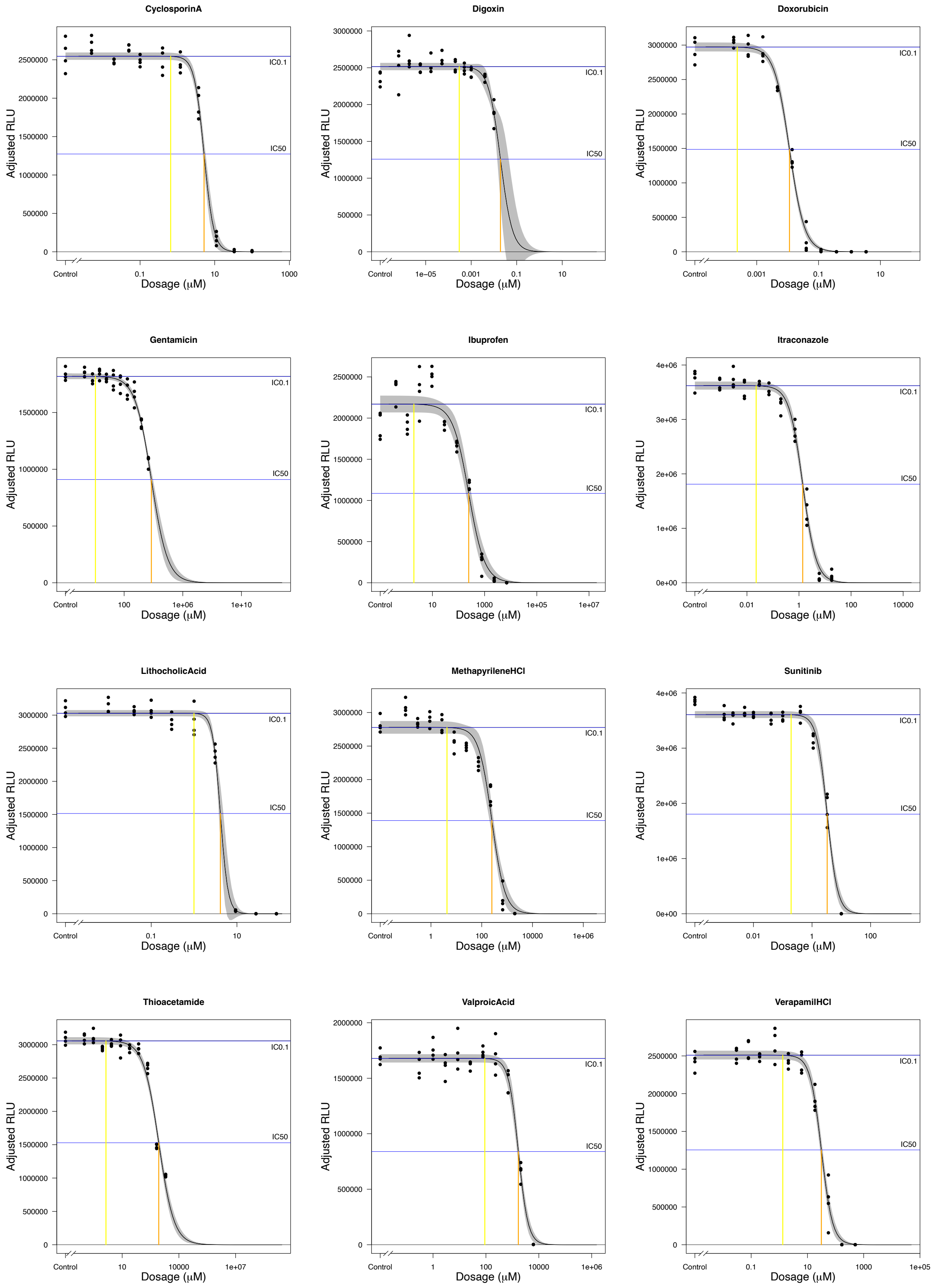


**Fig. S1 Dose response curves of the 24 chemicals in the ATP assay using KhES-3 cells**

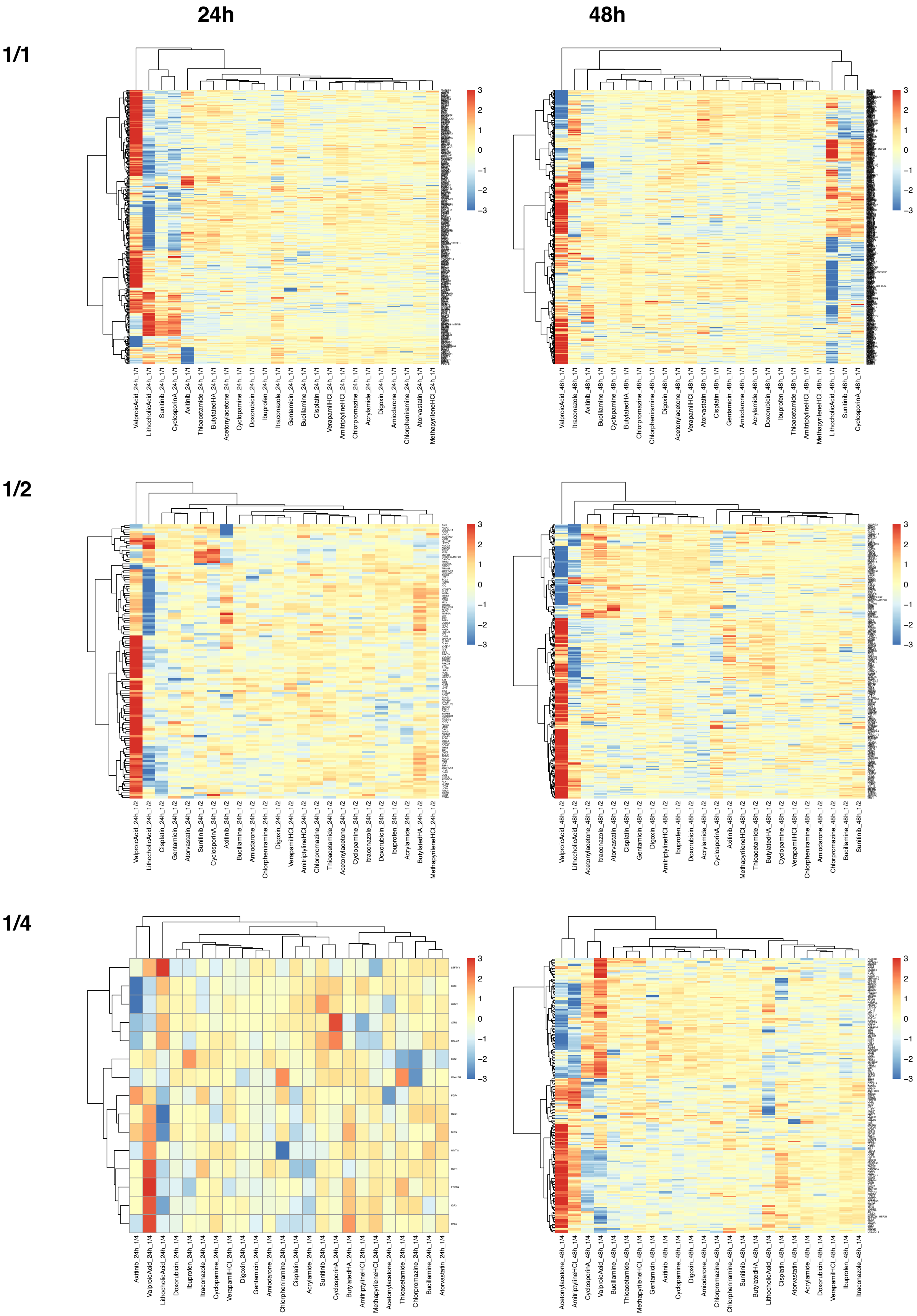
24h
48h
1/1
1/2
1/4
3
2
1
0
−1
−2
−3
ValproicAcid_24h_1/1
LithocholicAcid_24h_1/1
Sunitinib_24h_1/1
CyclosporinA_24h_1/1
Axitinib_24h_1/1
Thioacetamide_24h_1/1
ButylatedHA_24h_1/1
Acetonylacetone_24h_1/1
Cyclopamine_24h_1/1
Doxorubicin_24h_1/1
Ibuprofen_24h_1/1
Itraconazole_24h_1/1
Gentamicin_24h_1/1
Bucillamine_24h_1/1
Cisplatin_24h_1/1
VerapamilHCl_24h_1/1
AmitriptylineHCl_24h_1/1
Chlorpromazine_24h_1/1
Acrylamide_24h_1/1
Digoxin_24h_1/1
Amiodarone_24h_1/1
Chlorpheniramine_24h_1/1
Atorvastatin_24h_1/1
MethapyrileneHCl_24h_1/1
ValproicAcid_48h_1/1
Itraconazole_48h_1/1
Axitinib_48h_1/1
Bucillamine_48h_1/1
Cyclopamine_48h_1/1
ButylatedHA_48h_1/1
Chlorpromazine_48h_1/1
Chlorpheniramine_48h_1/1
Digoxin_48h_1/1
Acetonylacetone_48h_1/1
VerapamilHCl_48h_1/1
Atorvastatin_48h_1/1
Cisplatin_48h_1/1
Gentamicin_48h_1/1
Amiodarone_48h_1/1
Acrylamide_48h_1/1
Doxorubicin_48h_1/1
Ibuprofen_48h_1/1
Thioacetamide_48h_1/1
AmitriptylineHCl_48h_1/1
MethapyrileneHCl_48h_1/1
LithocholicAcid_48h_1/1
Sunitinib_48h_1/1
CyclosporinA_48h_1/1
ValproicAcid_24h_1/2
LithocholicAcid_24h_1/2
Cisplatin_24h_1/2
Gentamicin_24h_1/2
Atorvastatin_24h_1/2
Sunitinib_24h_1/2
CyclosporinA_24h_1/2
Axitinib_24h_1/2
Bucillamine_24h_1/2
Amiodarone_24h_1/2
Chlorpheniramine_24h_1/2
Digoxin_24h_1/2
VerapamilHCl_24h_1/2
AmitriptylineHCl_24h_1/2
Chlorpromazine_24h_1/2
Thioacetamide_24h_1/2
Acetonylacetone_24h_1/2
Cyclopamine_24h_1/2
Itraconazole_24h_1/2
Doxorubicin_24h_1/2
Ibuprofen_24h_1/2
Acrylamide_24h_1/2
ButylatedHA_24h_1/2
MethapyrileneHCl_24h_1/2
ValproicAcid_48h_1/2
LithocholicAcid_48h_1/2
Acetonylacetone_48h_1/2
Itraconazole_48h_1/2
Atorvastatin_48h_1/2
Cisplatin_48h_1/2
Gentamicin_48h_1/2
Digoxin_48h_1/2
AmitriptylineHCl_48h_1/2
Ibuprofen_48h_1/2
Doxorubicin_48h_1/2
Acrylamide_48h_1/2
CyclosporinA_48h_1/2
Axitinib_48h_1/2
MethapyrileneHCl_48h_1/2
Thioacetamide_48h_1/2
ButylatedHA_48h_1/2
Cyclopamine_48h_1/2
VerapamilHCl_48h_1/2
Chlorpheniramine_48h_1/2
Amiodarone_48h_1/2
Chlorpromazine_48h_1/2
Bucillamine_48h_1/2
Sunitinib_48h_1/2
Axitinib_24h_1/4
ValproicAcid_24h_1/4
LithocholicAcid_24h_1/4
Doxorubicin_24h_1/4
Ibuprofen_24h_1/4
Itraconazole_24h_1/4
Cyclopamine_24h_1/4
VerapamilHCl_24h_1/4
Digoxin_24h_1/4
Gentamicin_24h_1/4
Amiodarone_24h_1/4
Chlorpheniramine_24h_1/4
Cisplatin_24h_1/4
Acrylamide_24h_1/4
Sunitinib_24h_1/4
CyclosporinA_24h_1/4
ButylatedHA_24h_1/4
AmitriptylineHCl_24h_1/4
MethapyrileneHCl_24h_1/4
Acetonylacetone_24h_1/4
Thioacetamide_24h_1/4
Chlorpromazine_24h_1/4
Bucillamine_24h_1/4
Atorvastatin_24h_1/4
Acetonylacetone_48h_1/4
AmitriptylineHCl_48h_1/4
CyclosporinA_48h_1/4
ValproicAcid_48h_1/4
Bucillamine_48h_1/4
Thioacetamide_48h_1/4
MethapyrileneHCl_48h_1/4
Gentamicin_48h_1/4
Axitinib_48h_1/4
Cyclopamine_48h_1/4
Digoxin_48h_1/4
Amiodarone_48h_1/4
Chlorpromazine_48h_1/4
Chlorpheniramine_48h_1/4
Sunitinib_48h_1/4
ButylatedHA_48h_1/4
LithocholicAcid_48h_1/4
Cisplatin_48h_1/4
Atorvastatin_48h_1/4
Acrylamide_48h_1/4
Doxorubicin_48h_1/4
VerapamilHCl_48h_1/4
Ibuprofen_48h_1/4
Itraconazole_48h_1/4

**1/8**

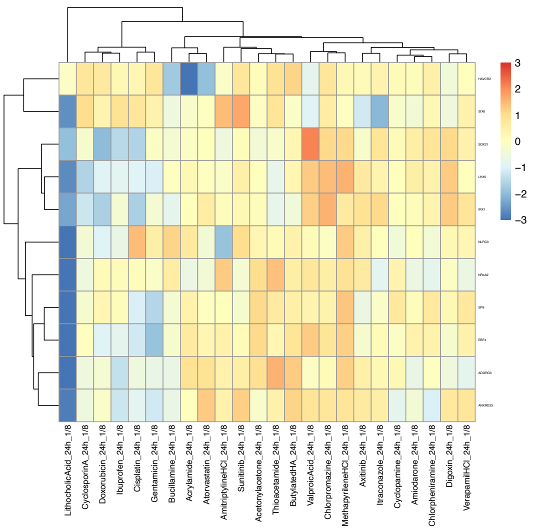


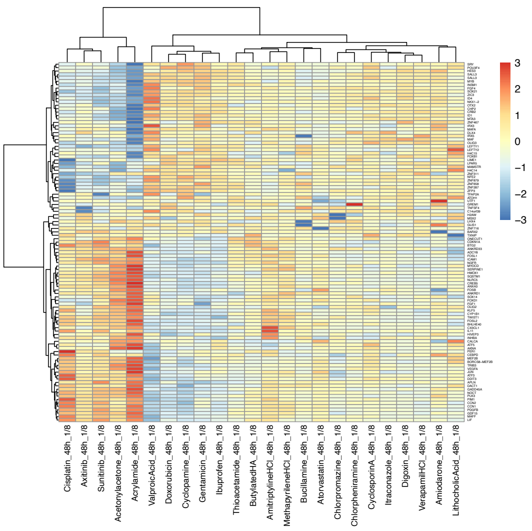


**1/16**

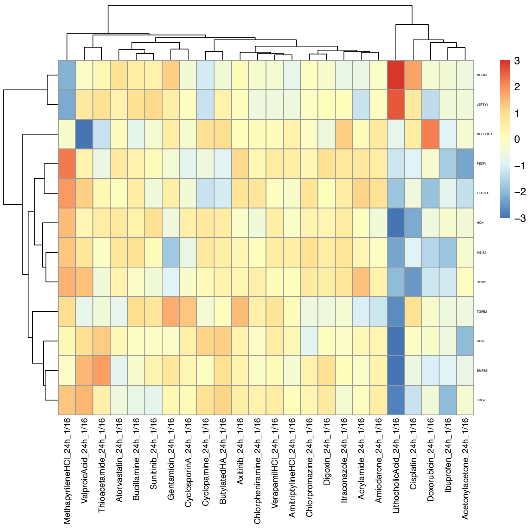


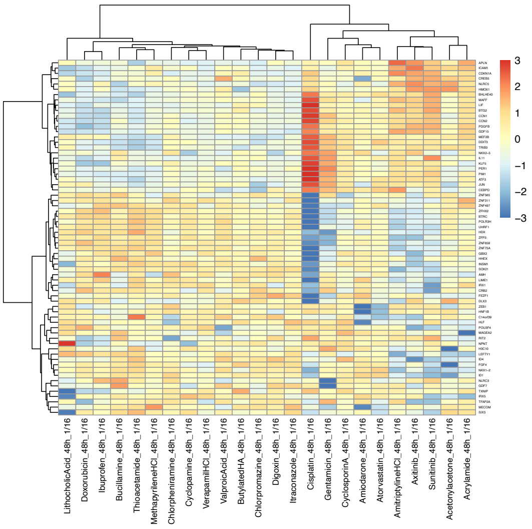


**Fig. S2 Differentially expressed transcription factor genes for 24 chemicals at five doses at 24 h and 48 h**

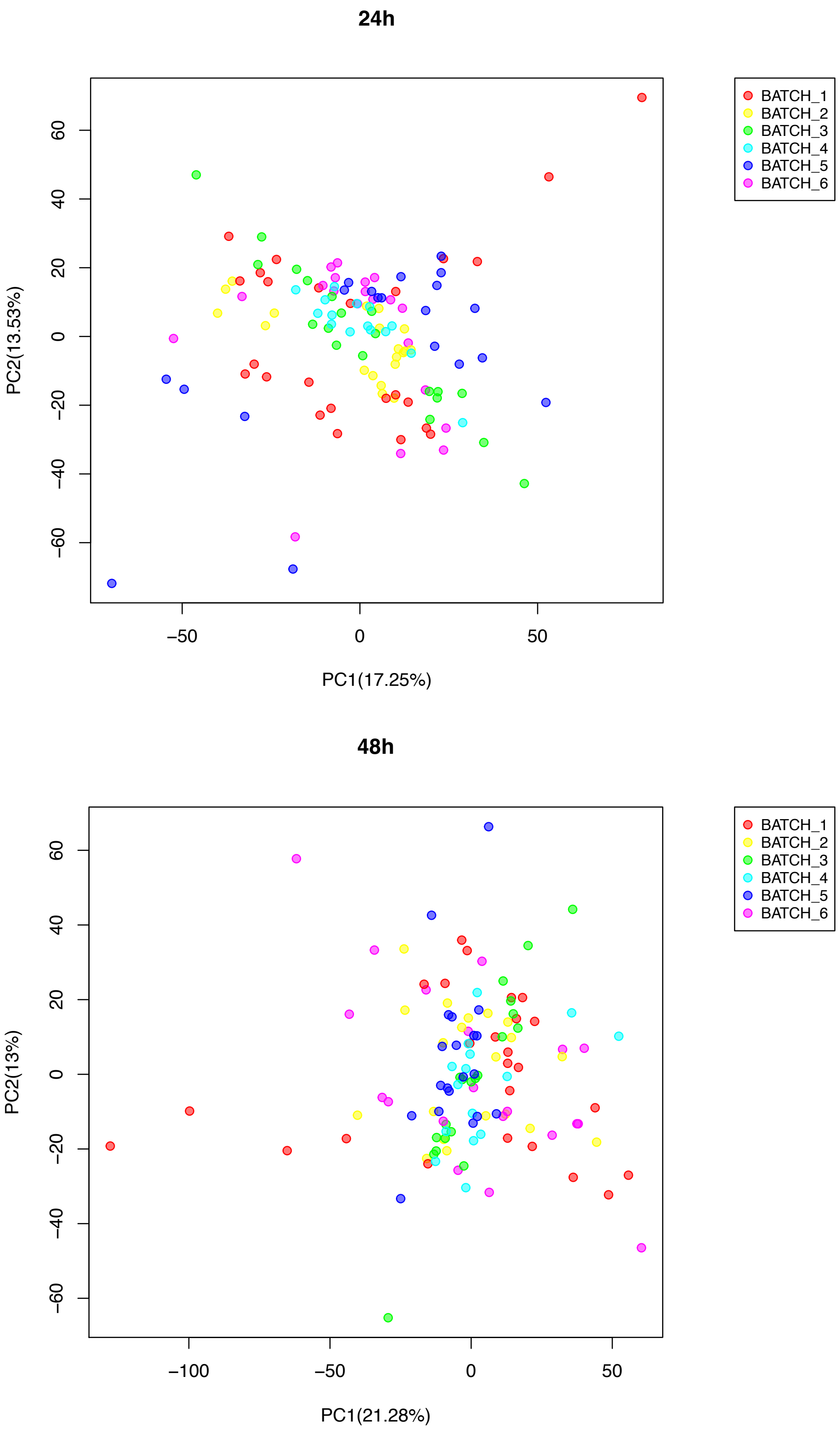


**Fig. S3 PCA analysis of KhES-3 cell data at 24 h and 48 h**

#### Acetonylacetone

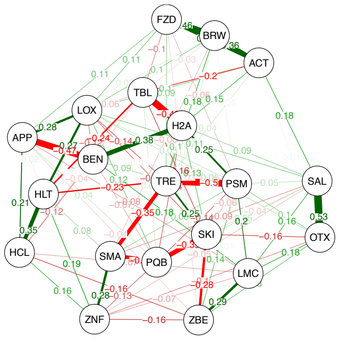


#### Acrylamide

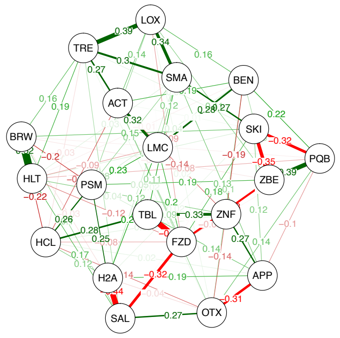


#### Amiodarone

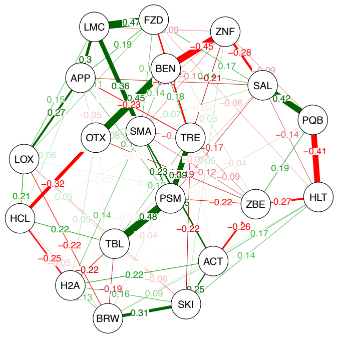


#### AmitriptylineHCl

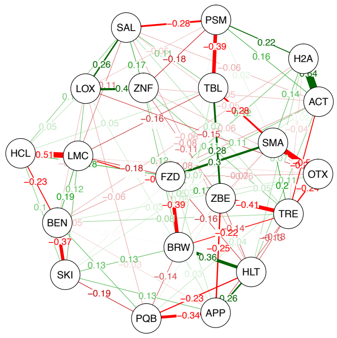


#### Atorvastatin

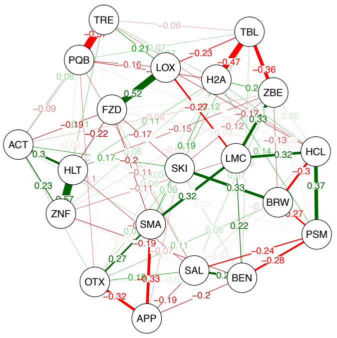


#### Axitinib

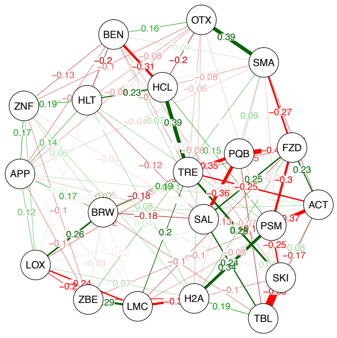


#### Bucillamine

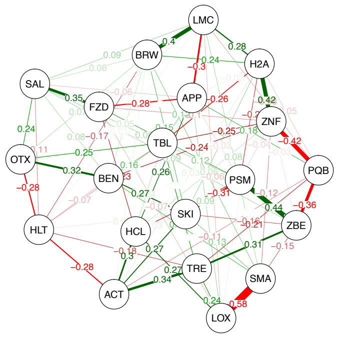


#### ButylatedHA

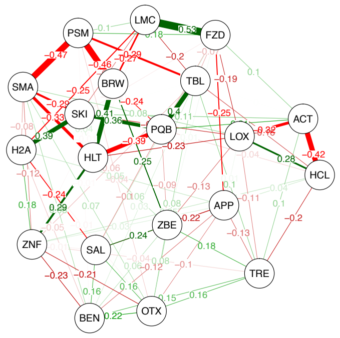


#### Chlorpheniramine

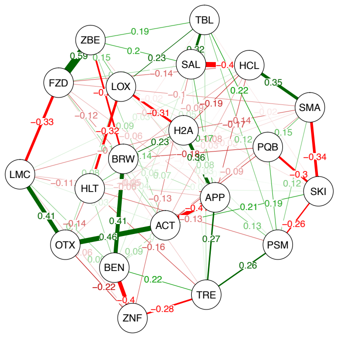


#### Chlorpromazine

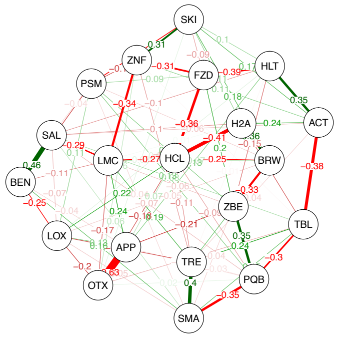


#### Cisplatin

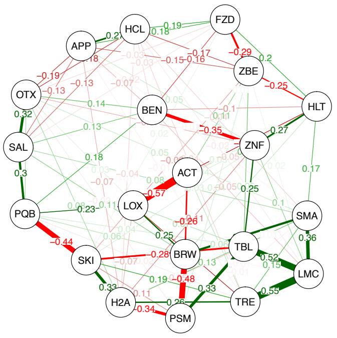


#### Cyclopamine

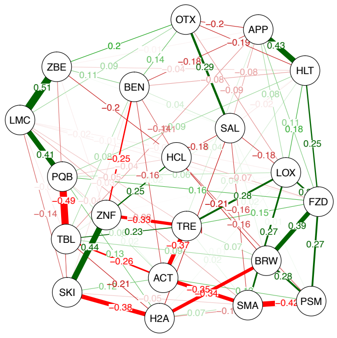

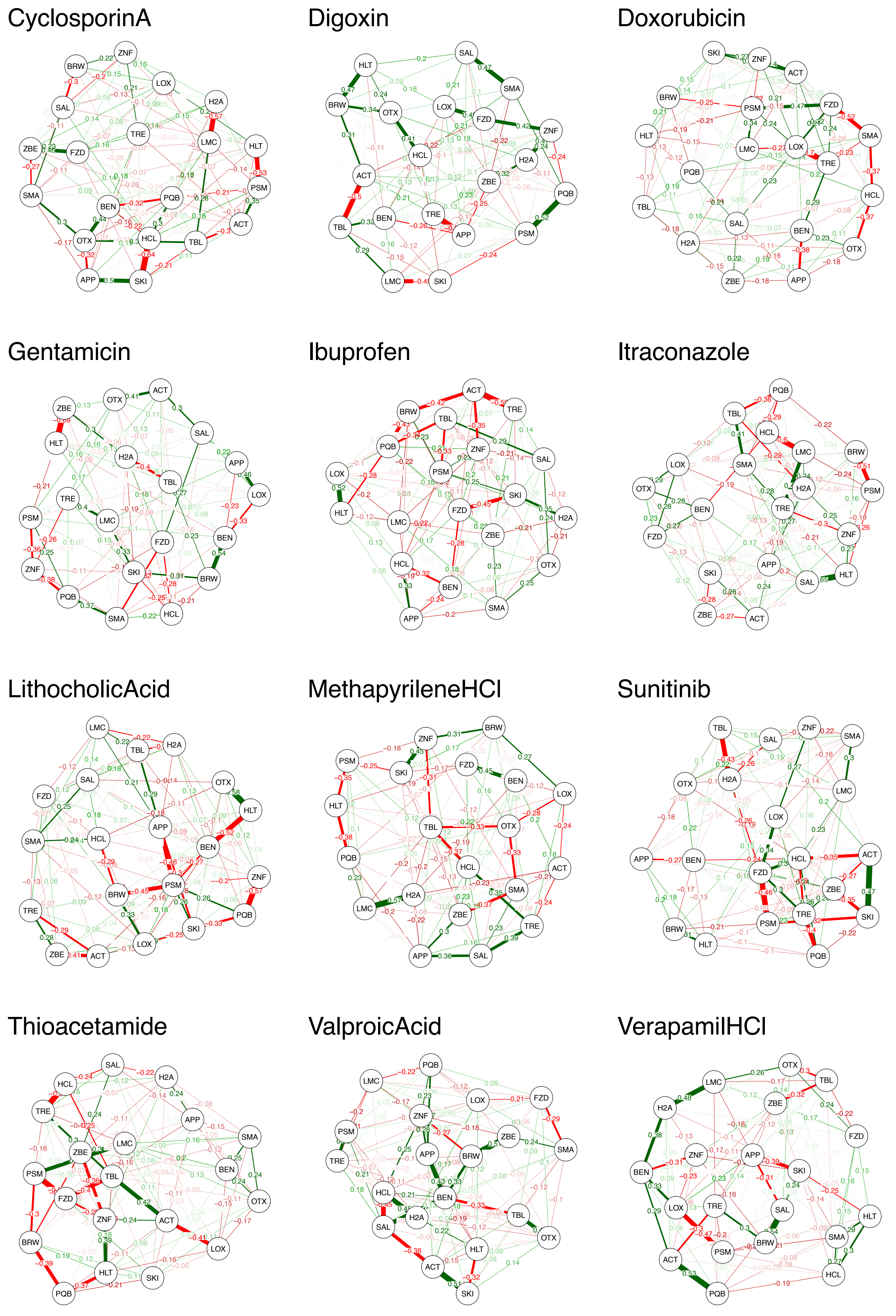


**Fig. S4 GGM for each of the 24 chemicals at 24 h**

Acetonylacetone

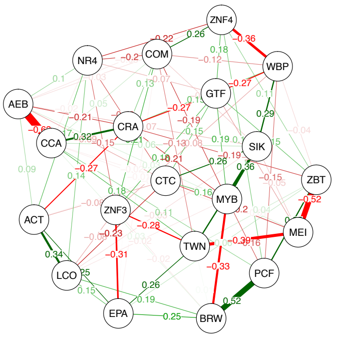

Acrylamide

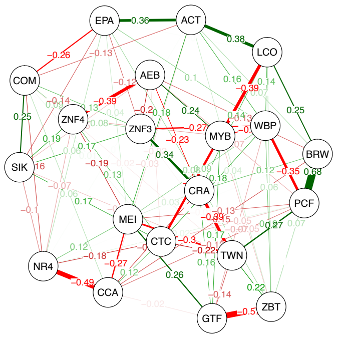

Amiodarone

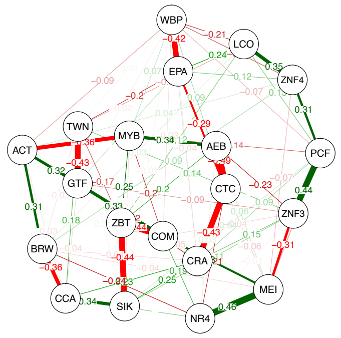

AmitriptylineHCl

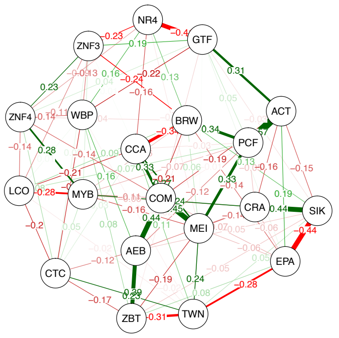

Atorvastatin

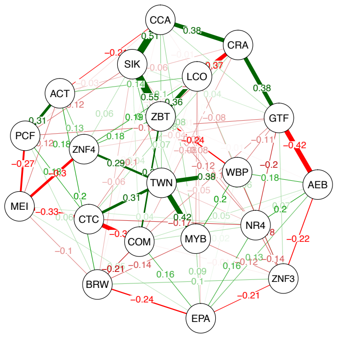

Axitinib

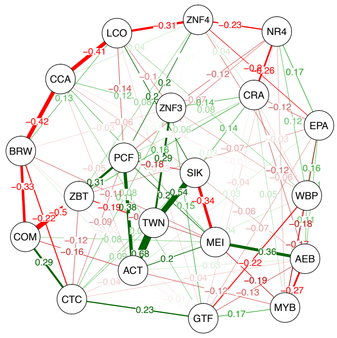

Bucillamine

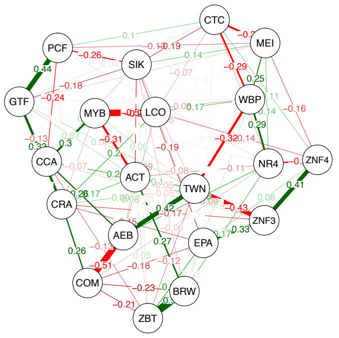

ButylatedHA

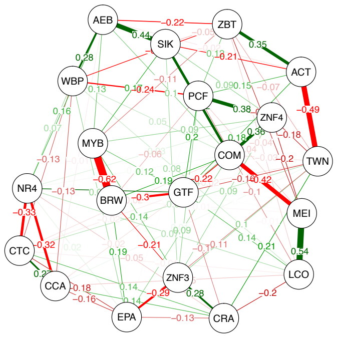

Chlorpheniramine

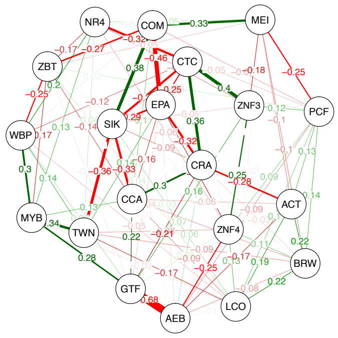

Chlorpromazine

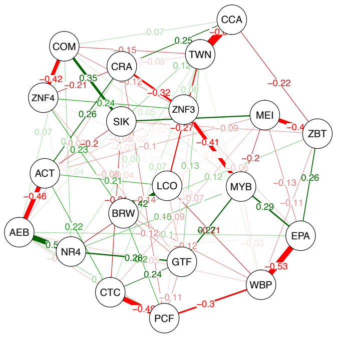

Cisplatin

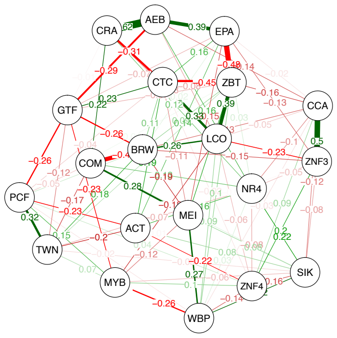

Cyclopamine

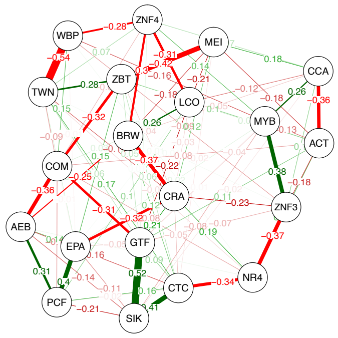

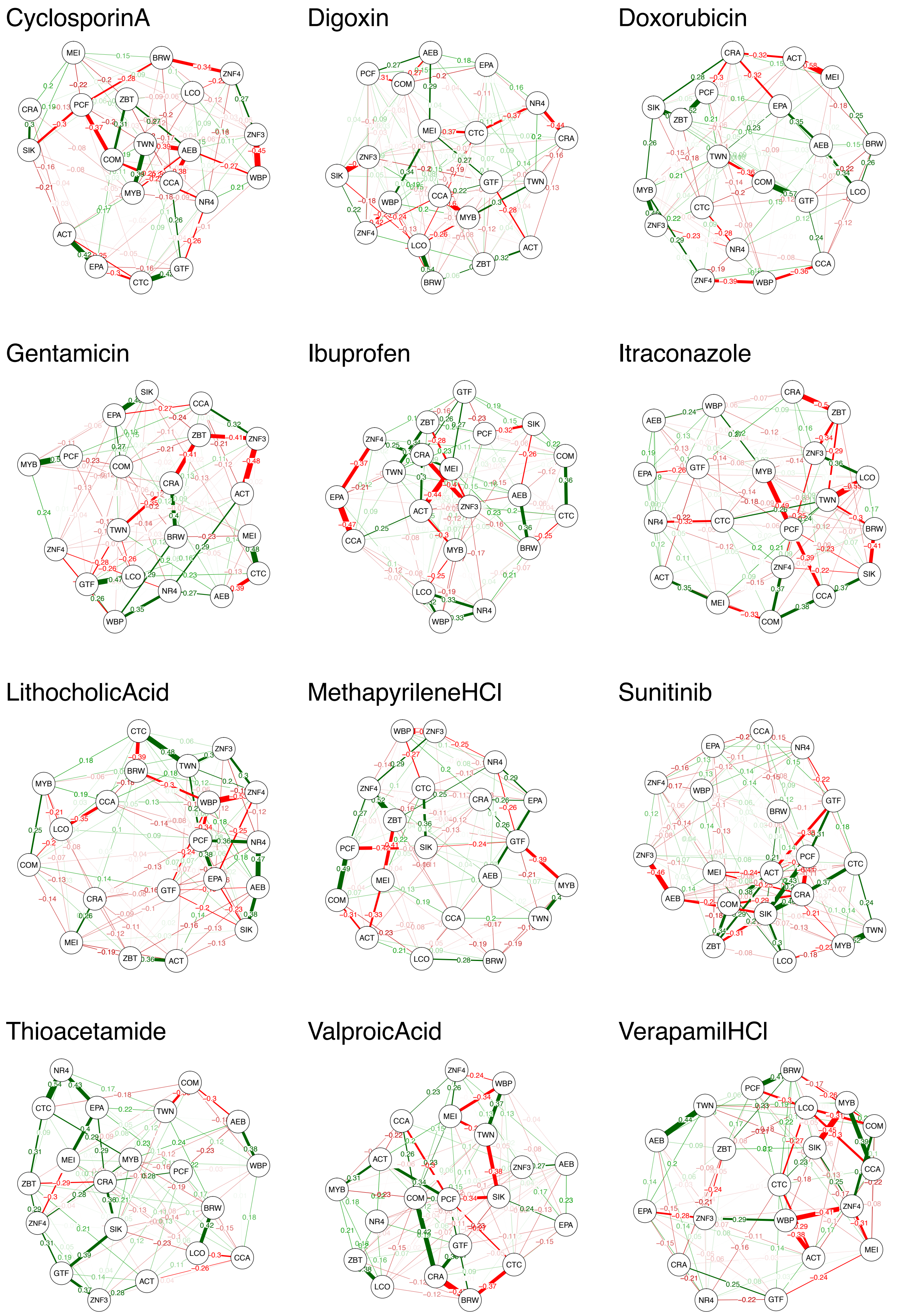


**Fig. S5 GGM for each of the 24 chemicals at 48 h**

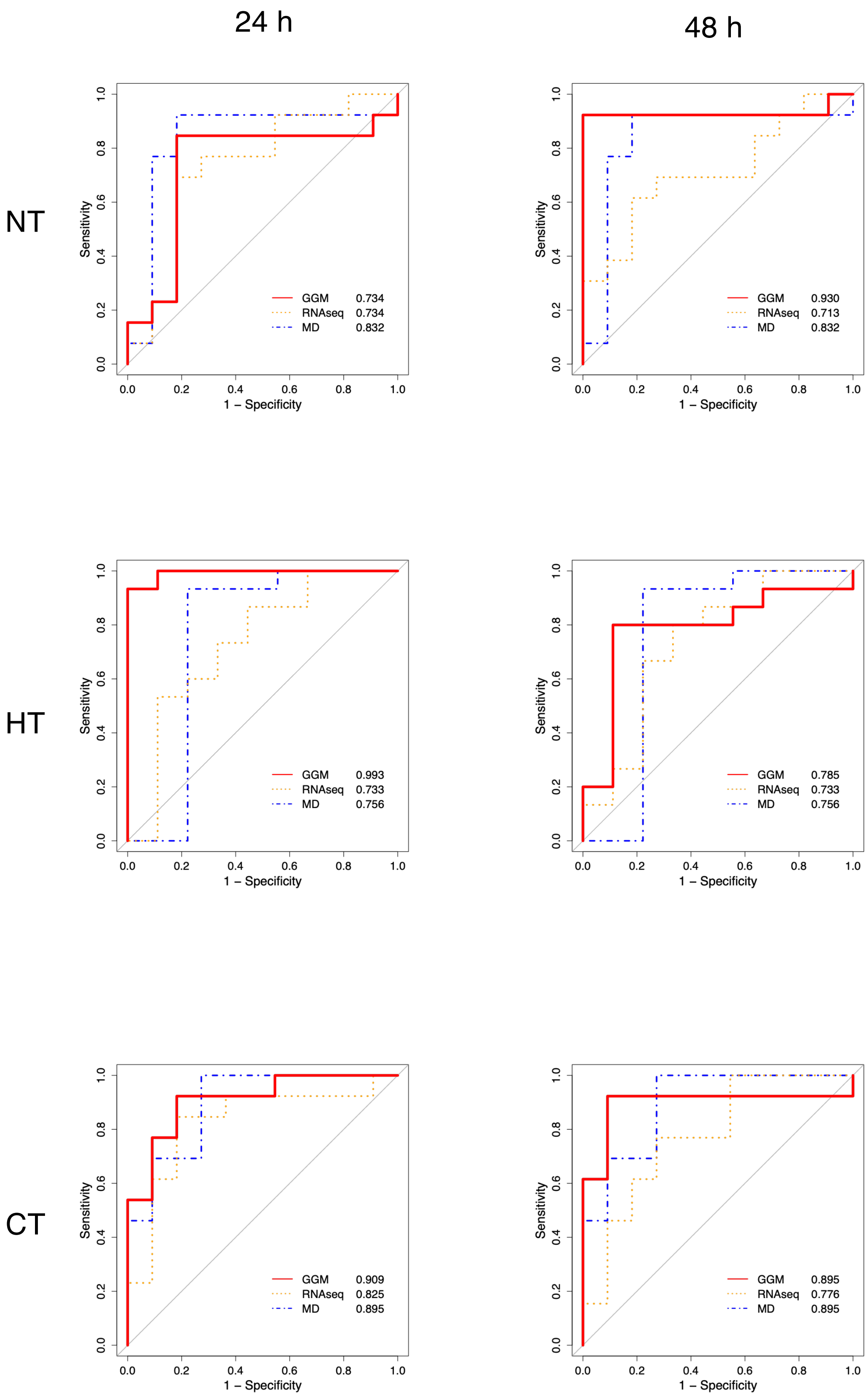
24 h
48 h
NT
HT
CT
Sensitivity
1 – Specificity
GGM 0.734
RNAseq 0.734
MD 0.832
GGM 0.930
RNAseq 0.713
MD 0.832
GGM 0.993
RNAseq 0.733
MD 0.756
GGM 0.785
RNAseq 0.733
MD 0.756
GGM 0.909
RNAseq 0.825
MD 0.895
GGM 0.895
RNAseq 0.776
MD 0.895

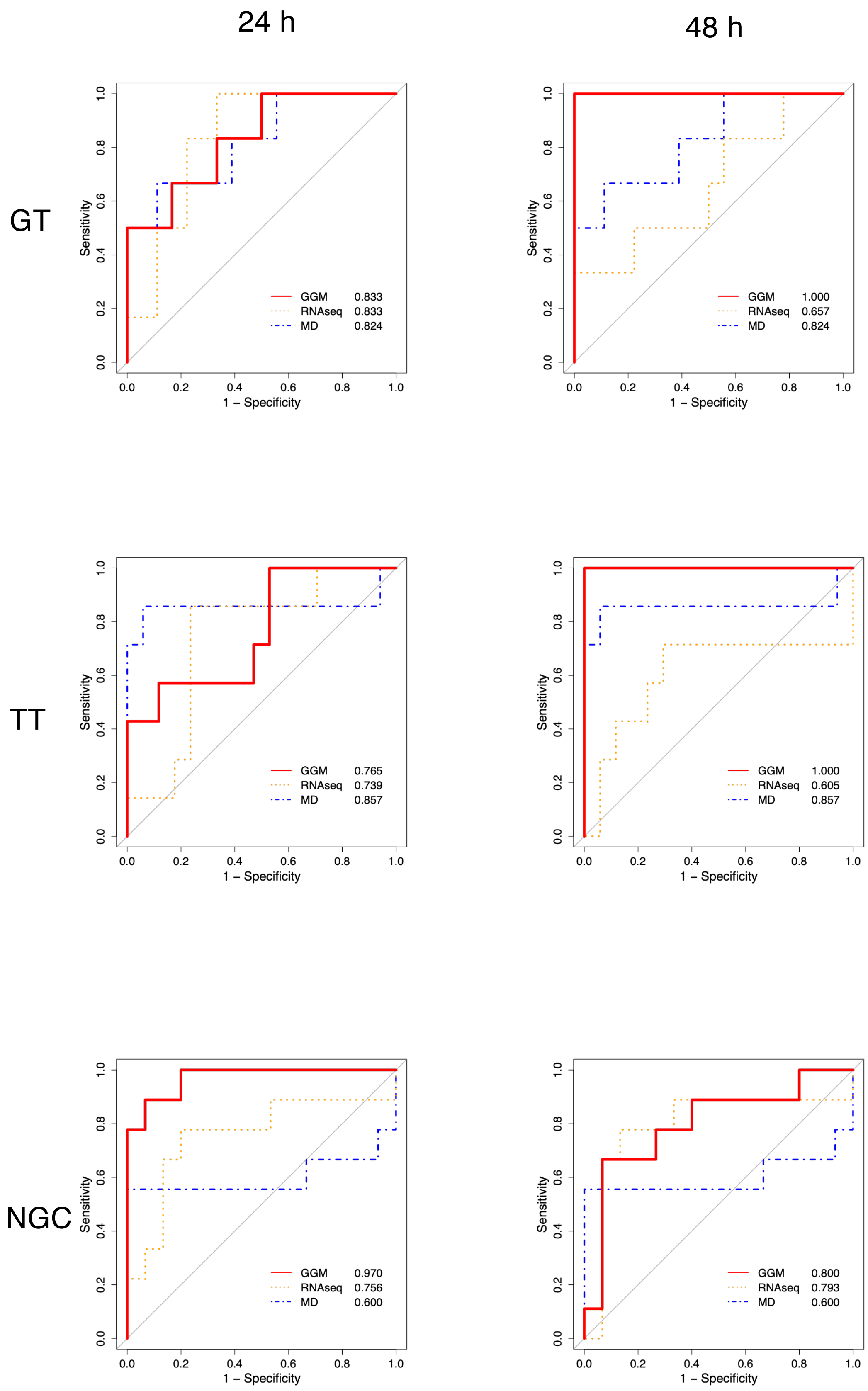


**Fig. S6 ROC curves for predictions of 6 toxic categories at 24 h and 48 h**

# NT 24h

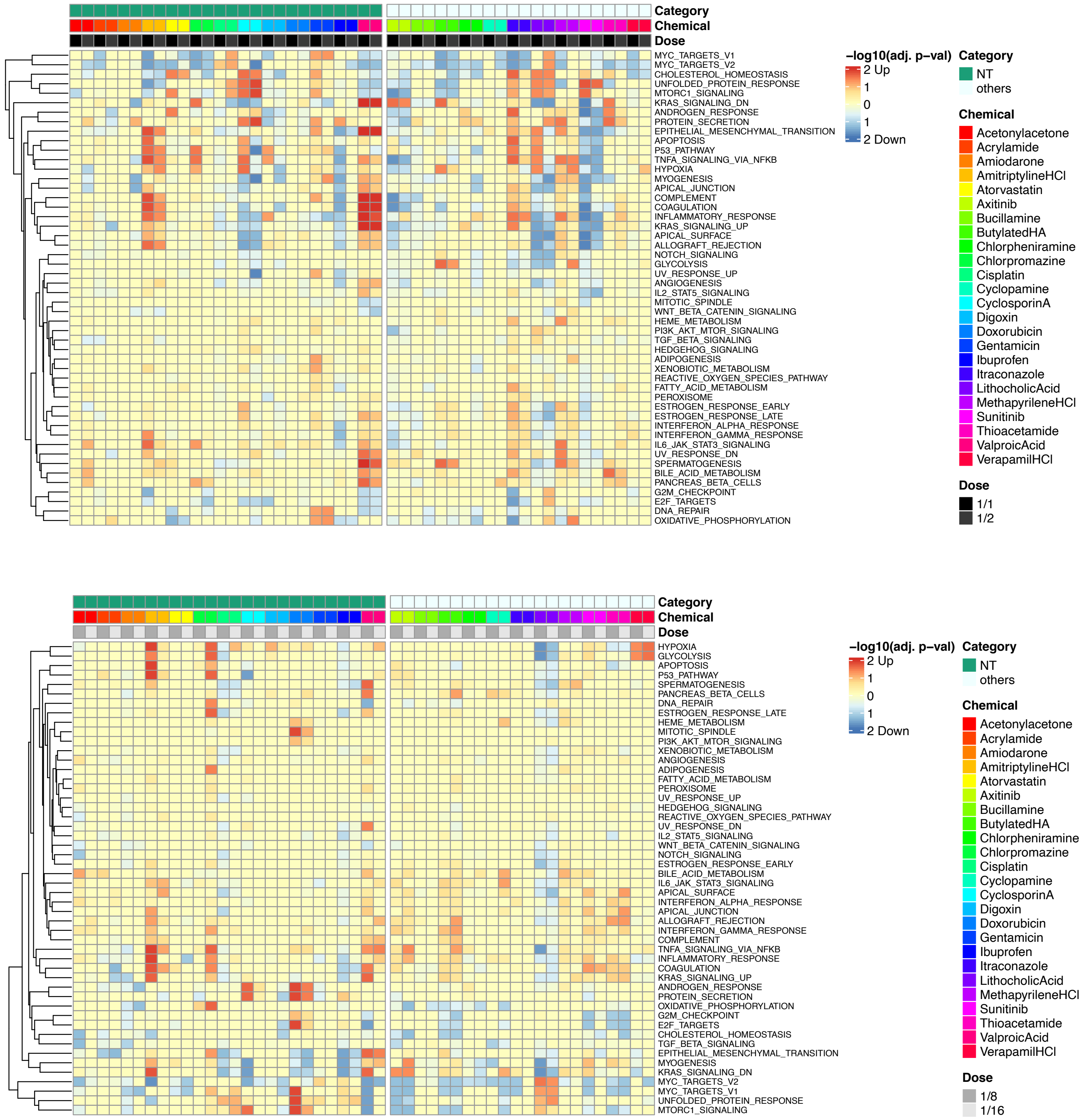


upper: high doses
lower: low doses

# HT 24h

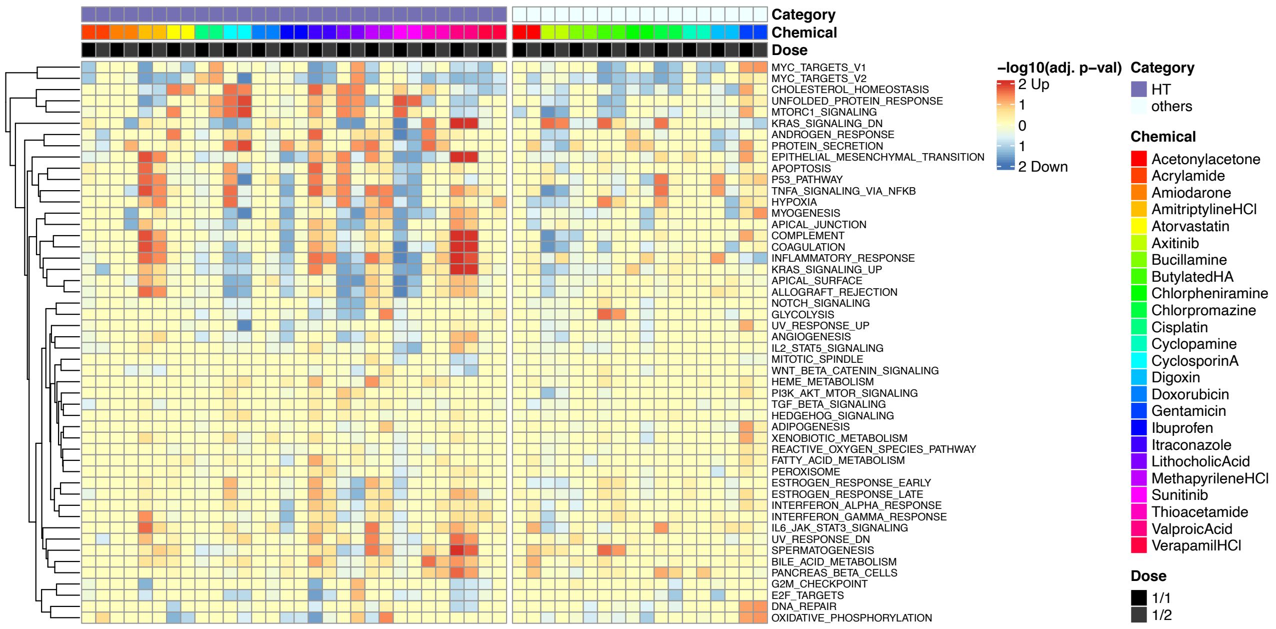


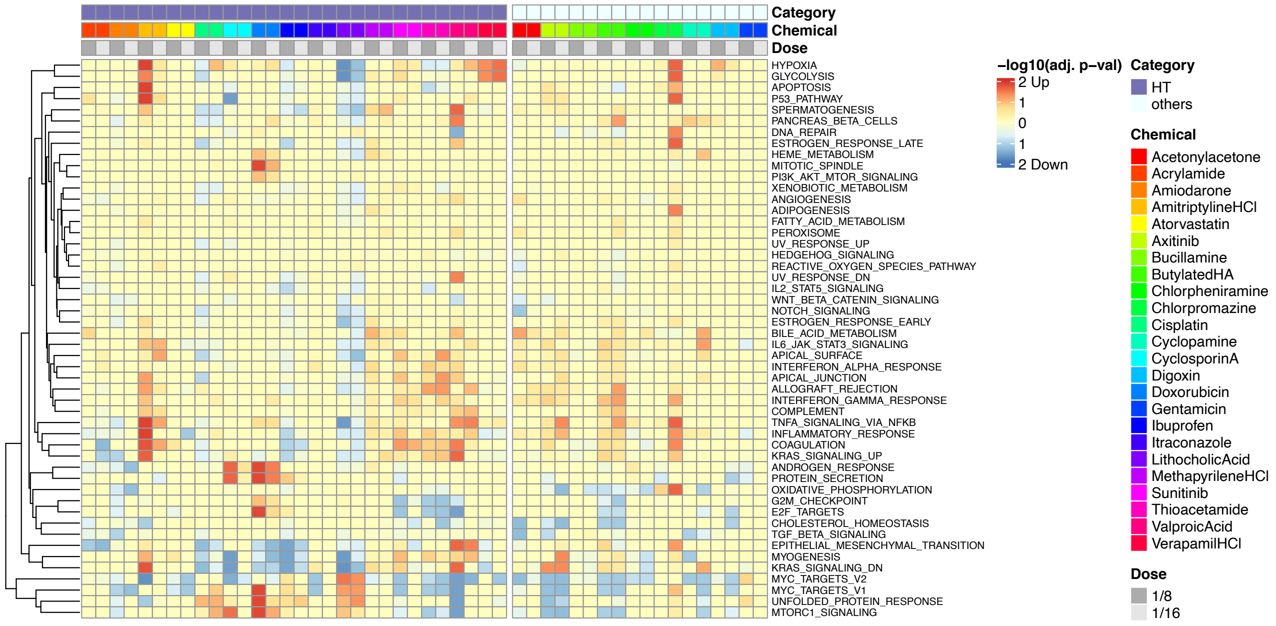


upper: high doses

lower: low doses

# CT 24h

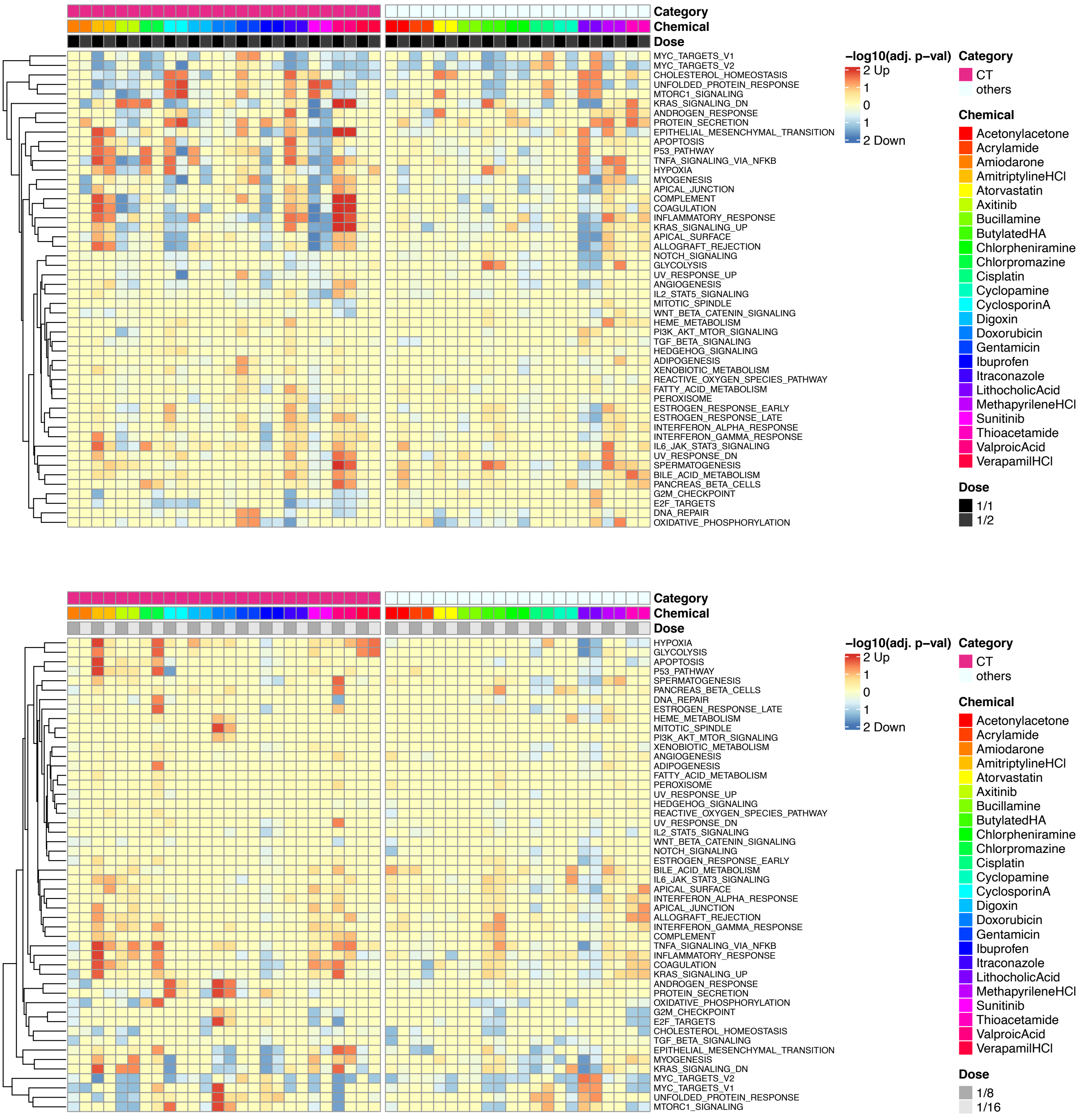


upper: high doses
lower: low doses

# GT 24h

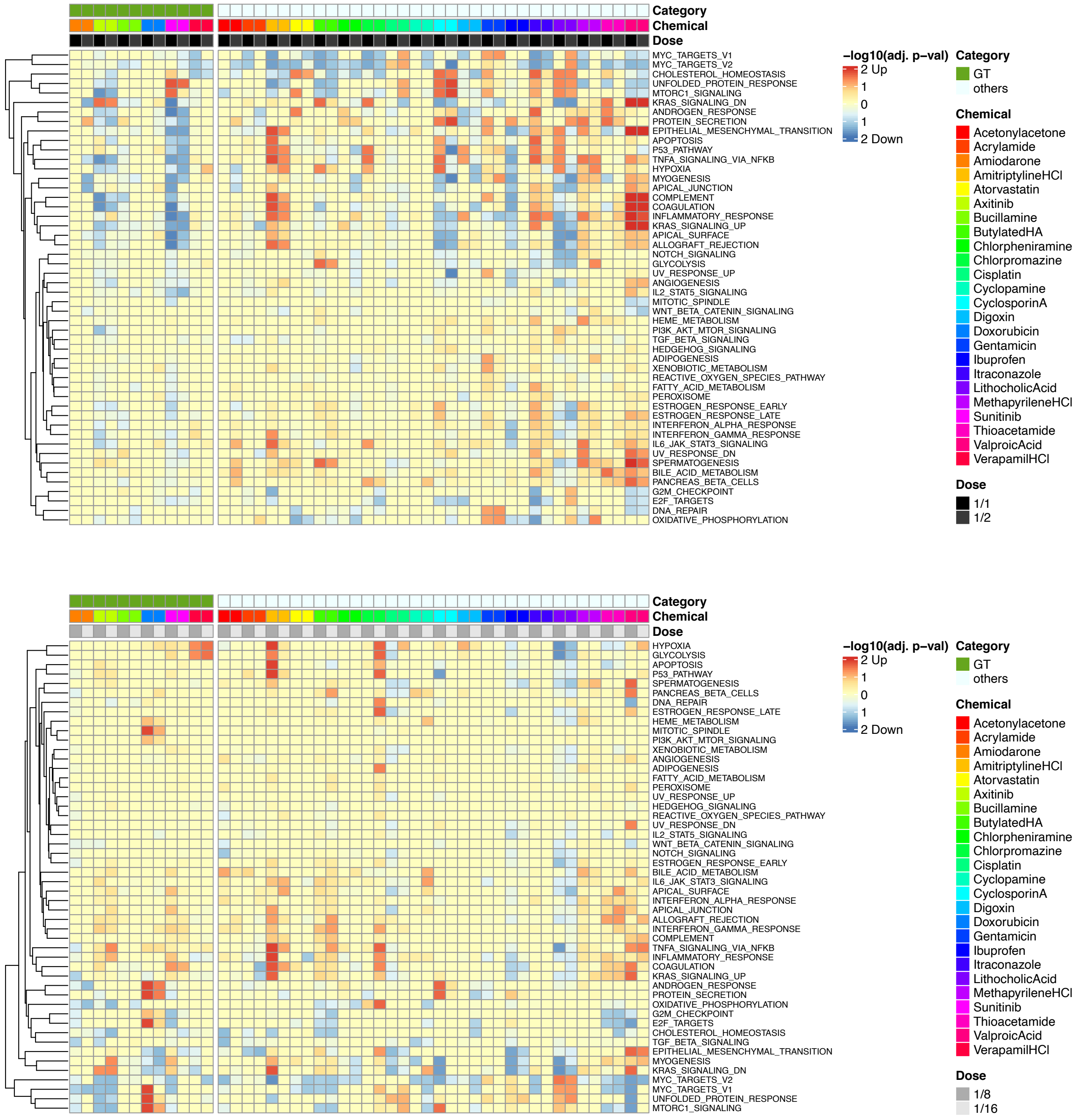


upper: high doses
lower: low doses

# TT 24h

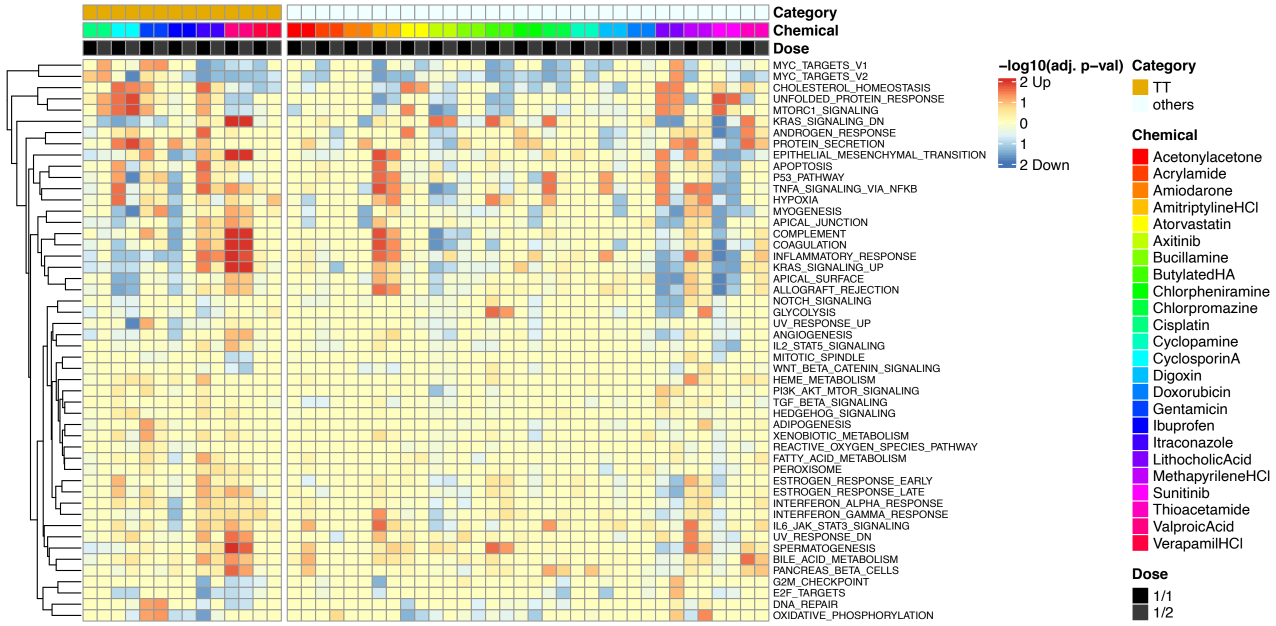


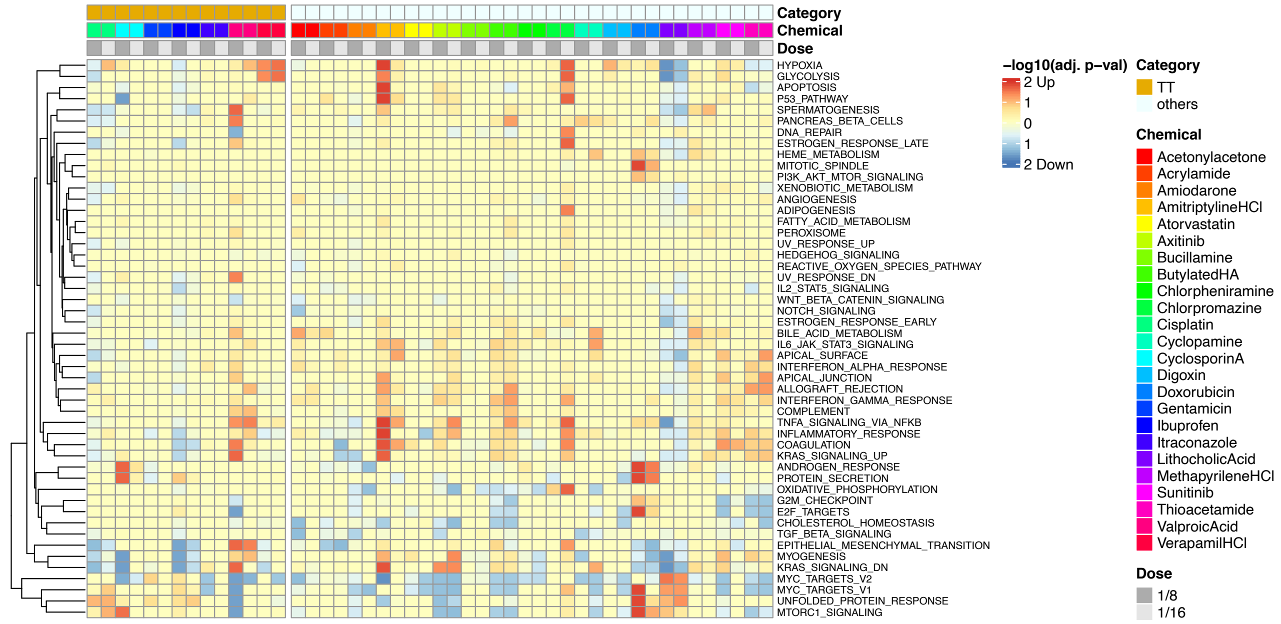


upper: high doses
lower: low doses

### NGC 24h

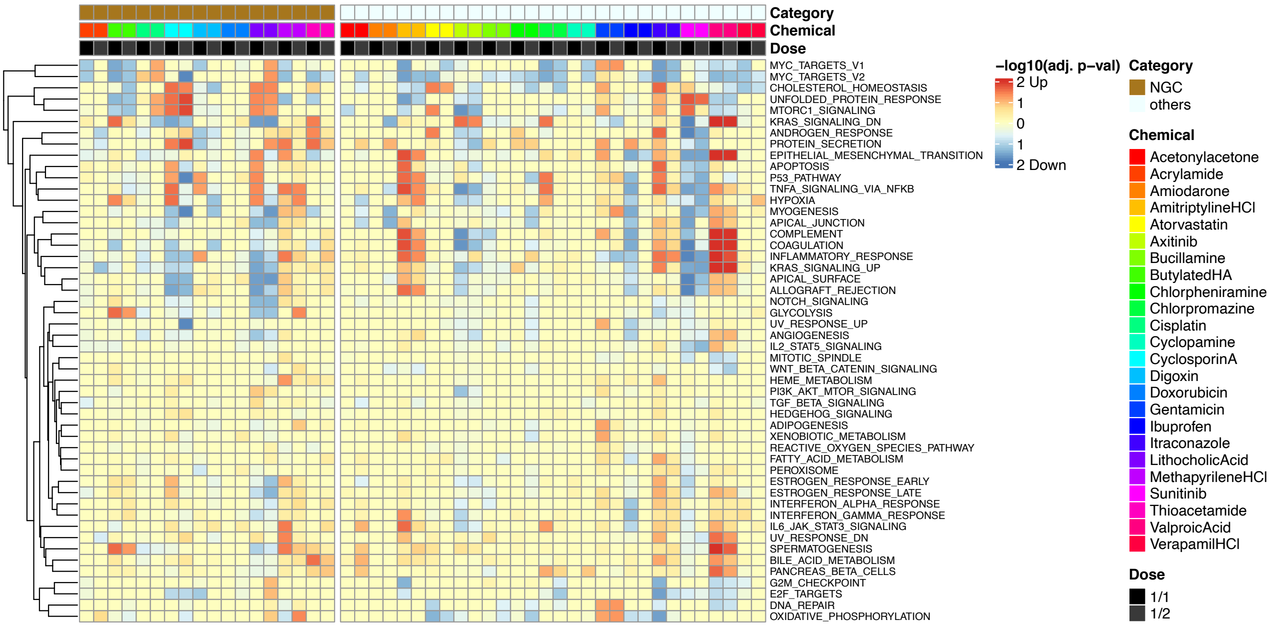


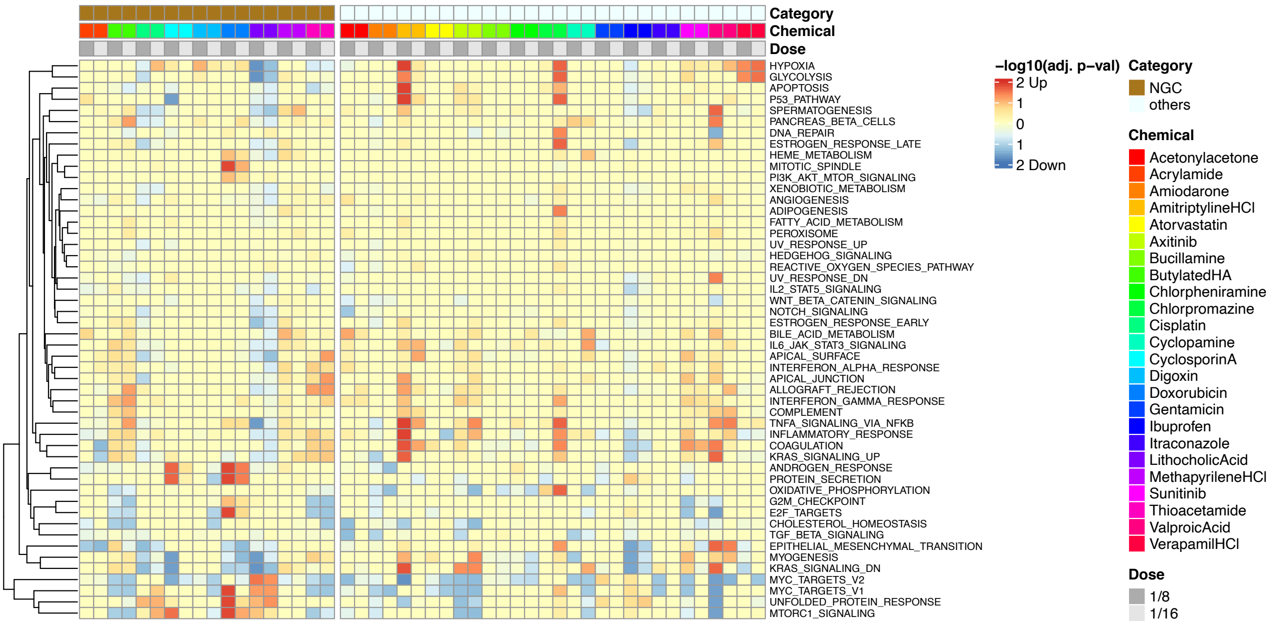


upper: high doses
lower: low doses

# NT 48h

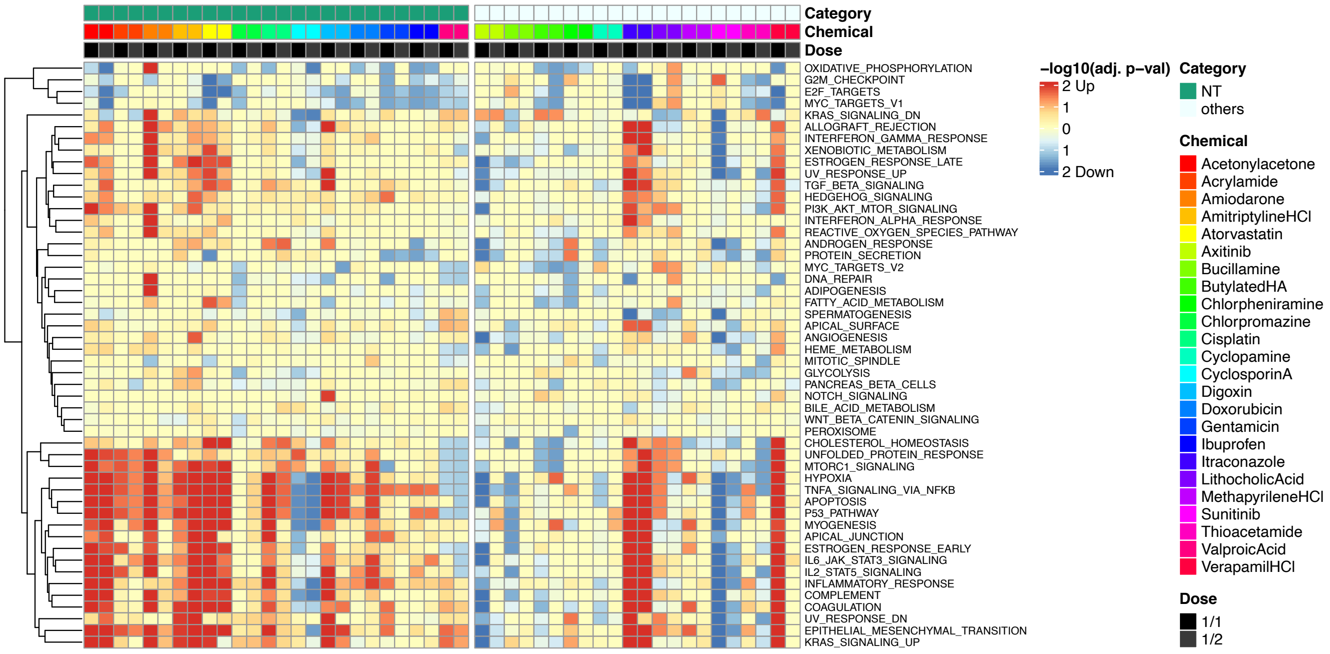


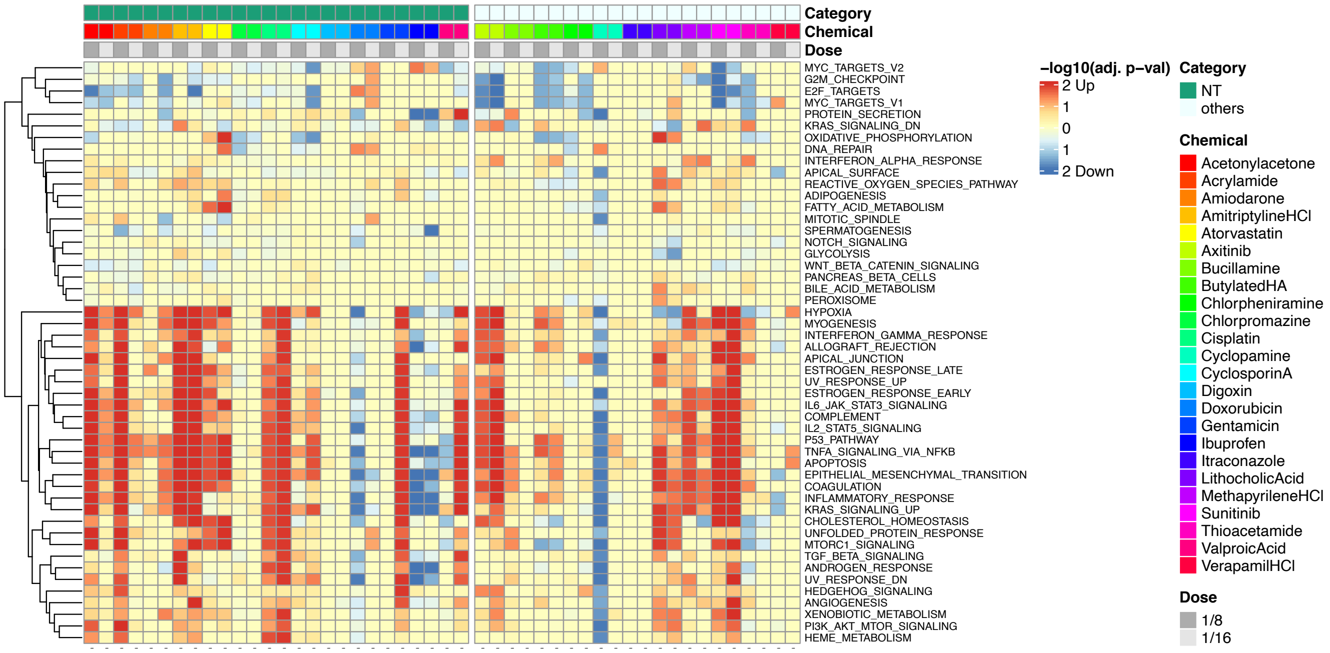


upper: high doses
lower: low doses

# HT 48h

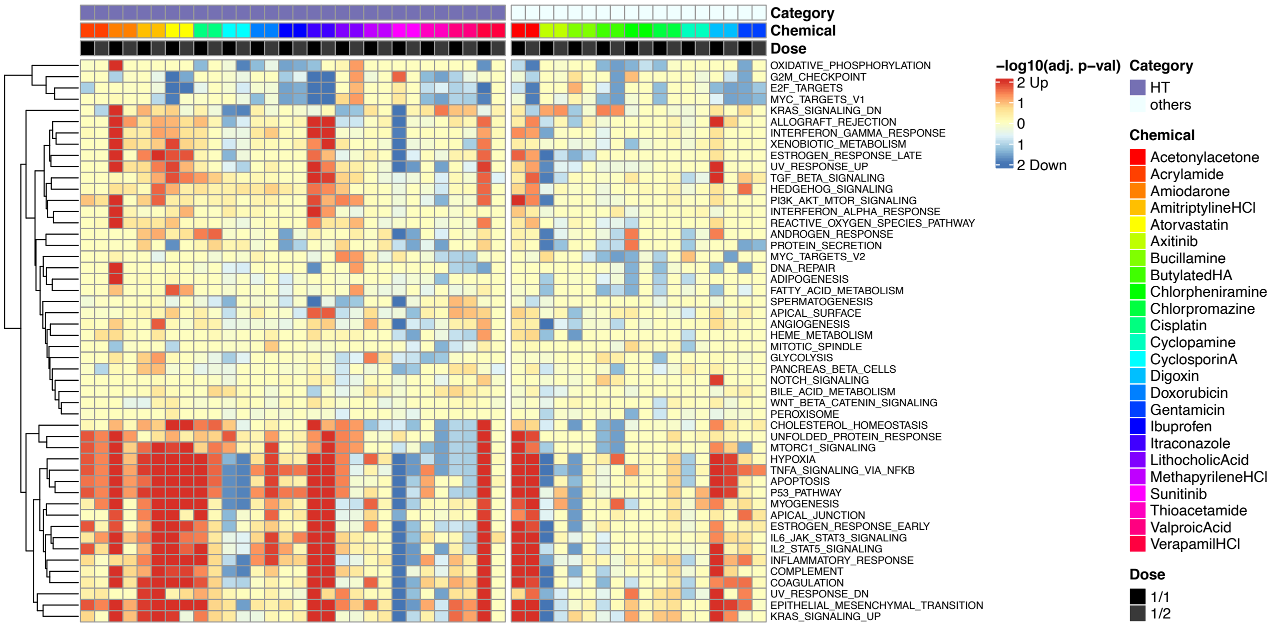


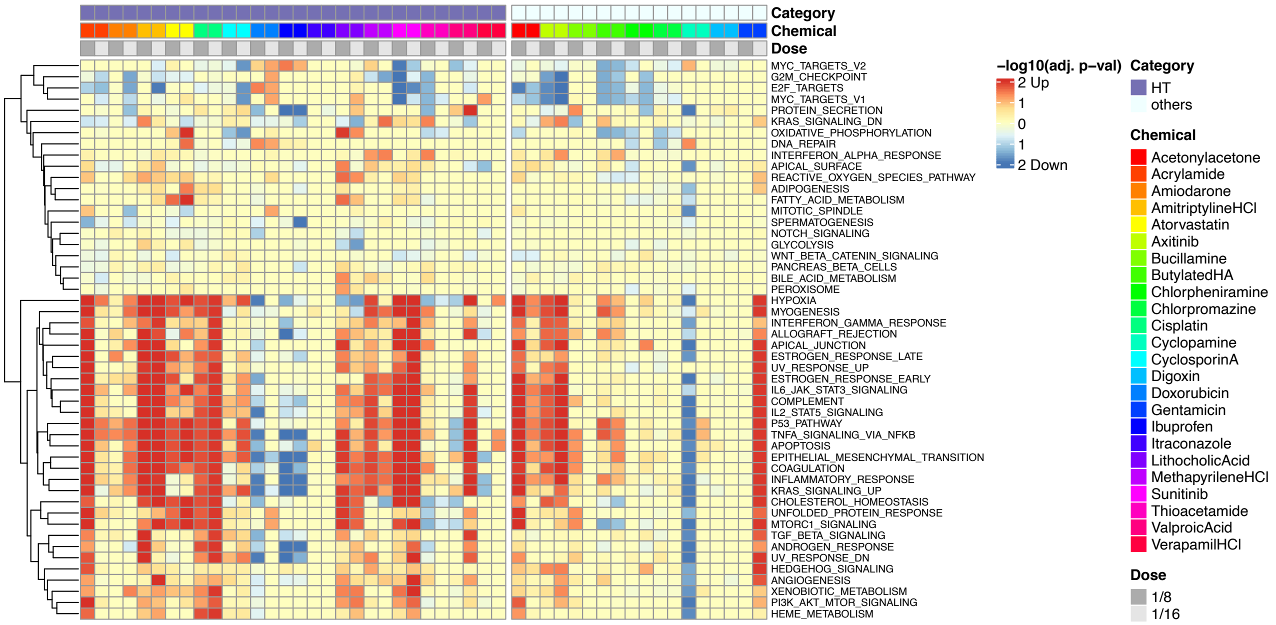


upper: high doses
lower: low doses

# CT 48h

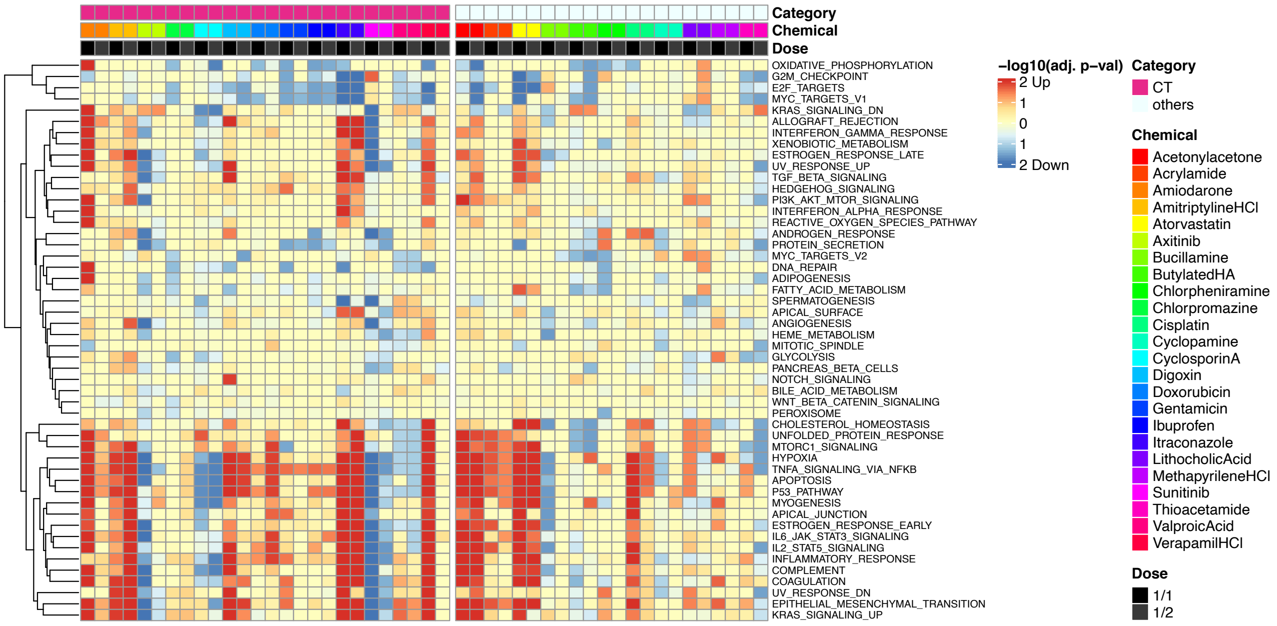


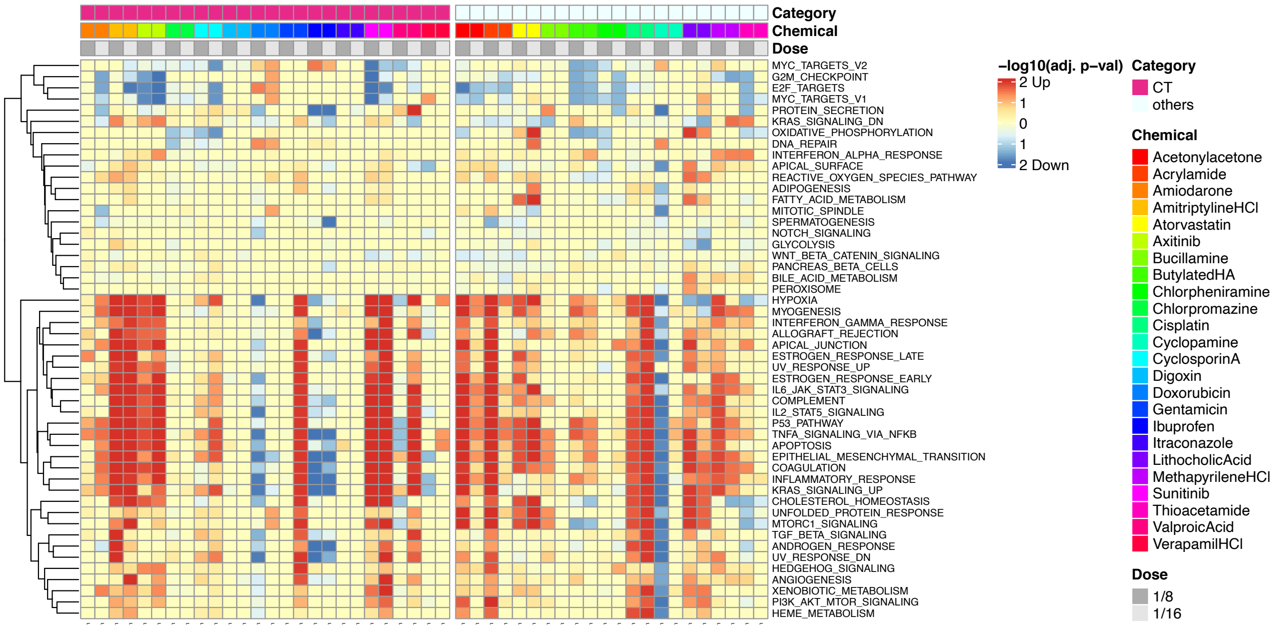


upper: high doses
lower: low doses

# GT 48h

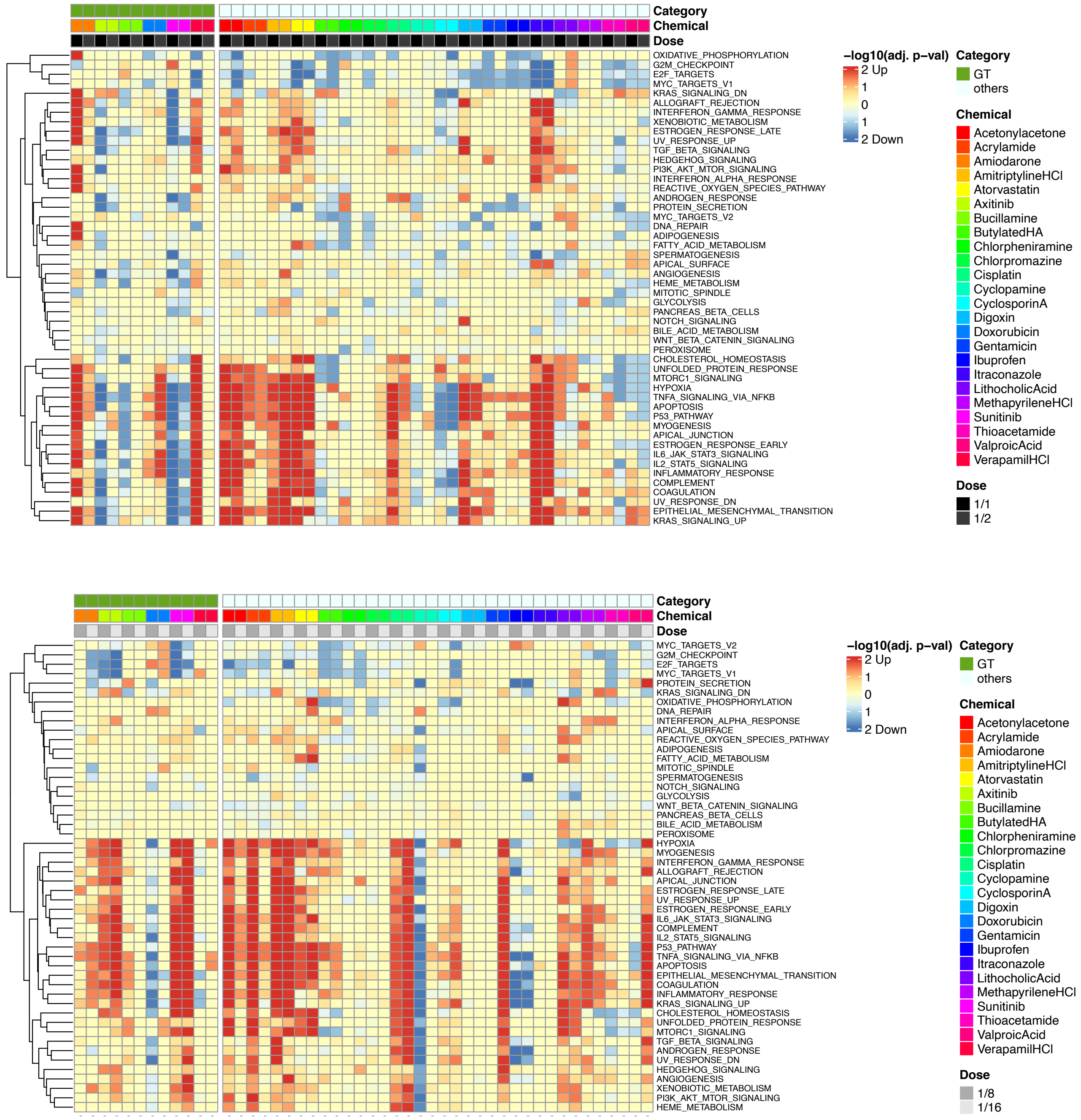


upper: high doses
lower: low doses

# TT 48h

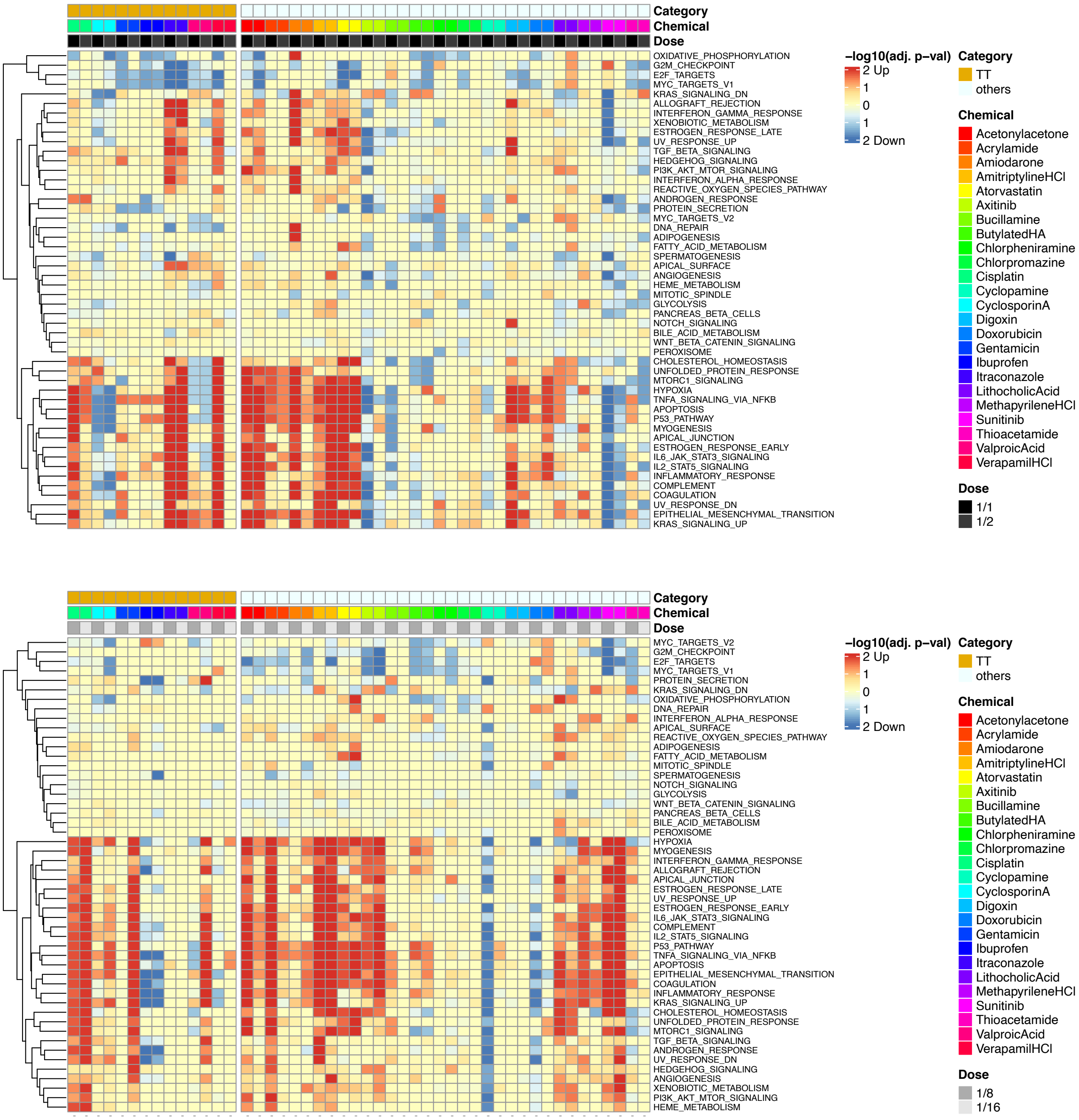


upper: high doses
lower: low doses

### NGC 48h

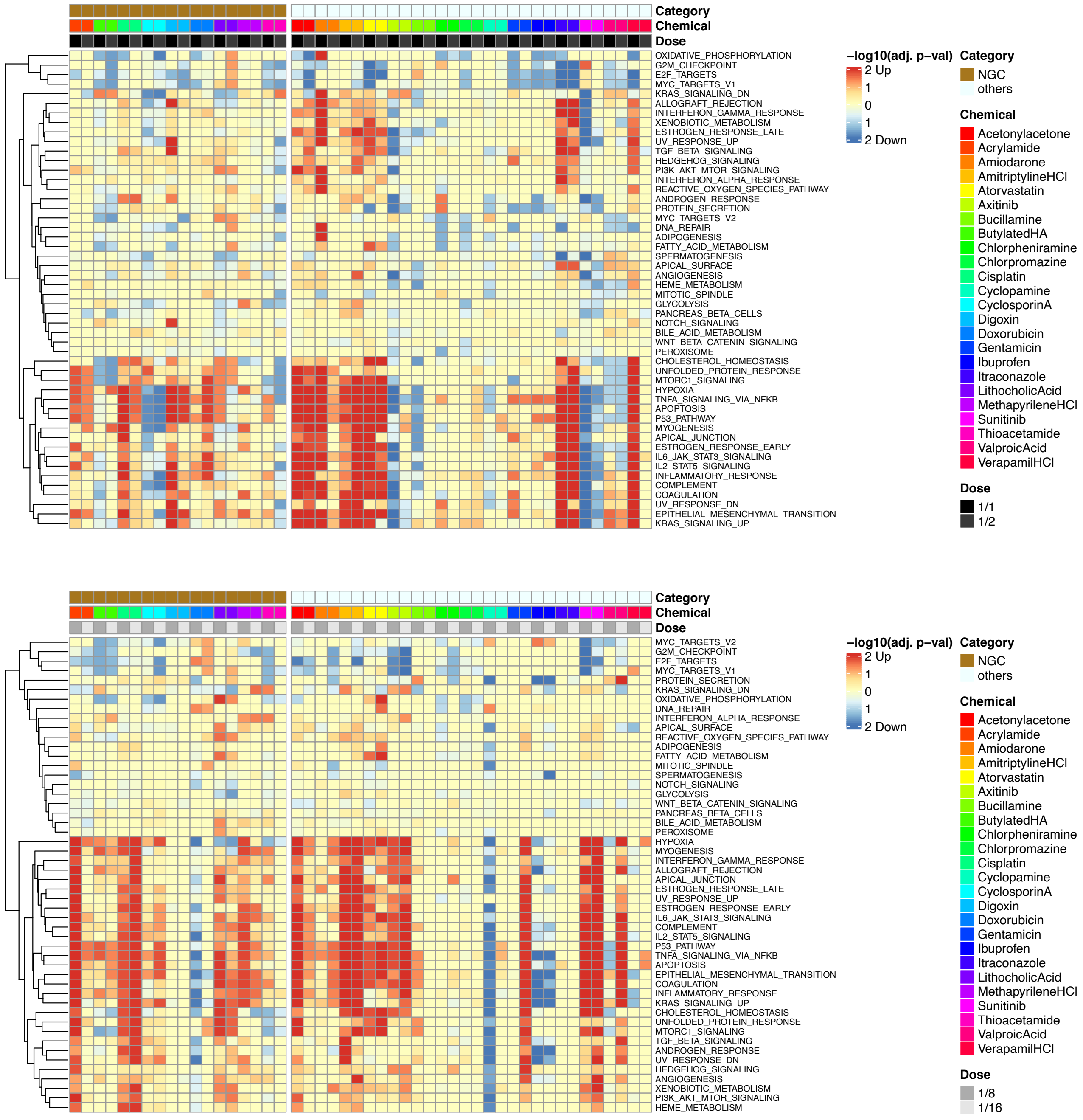


upper: high doses
lower: low doses

**Fig. S7 Pathway analysis by GSEA of the 6 toxic categories**

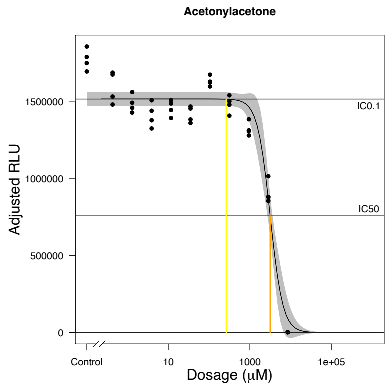
Acetonylacetone
1500000
1000000
500000
0
Adjusted RLU
IC0.1
IC50
Control
10
1000
1e+05
Dosage (μM)

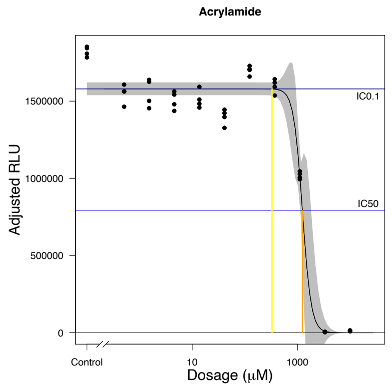
Acrylamide
1500000
1000000
500000
0
Adjusted RLU
IC0.1
IC50
Control
10
1000
Dosage (μM)

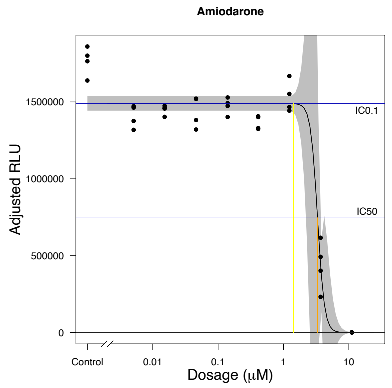
Amiodarone
1500000
1000000
500000
0
Adjusted RLU
IC0.1
IC50
Control
0.01
0.1
1
10
Dosage (μM)

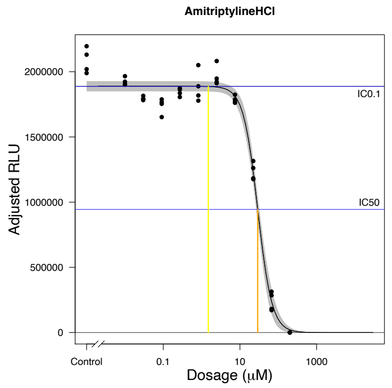
AmitriptylineHCl
2000000
1500000
1000000
500000
0
Adjusted RLU
IC0.1
IC50
Control
0.1
10
1000
Dosage (μM)

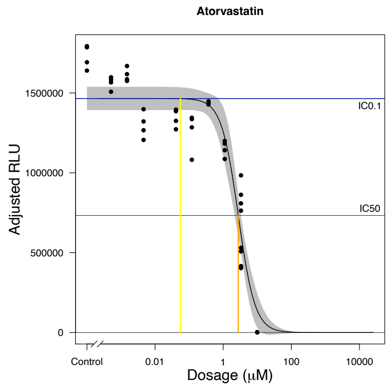
Atorvastatin
1500000
1000000
500000
0
Adjusted RLU
IC0.1
IC50
Control
0.01
1
100
10000
Dosage (μM)

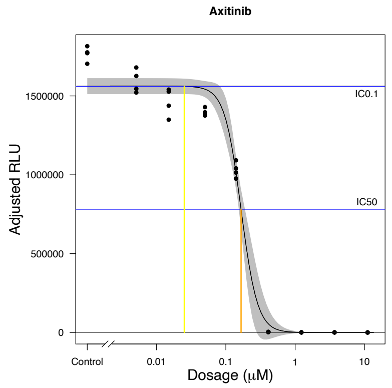
Axitinib
1500000
1000000
500000
0
Adjusted RLU
IC0.1
IC50
Control
0.01
0.1
1
10
Dosage (μM)

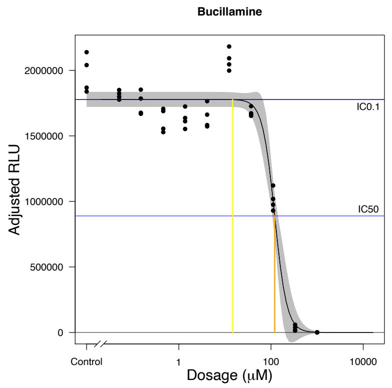
Bucillamine
2000000
1500000
1000000
500000
0
Adjusted RLU
IC0.1
IC50
Control
1
100
10000
Dosage (μM)

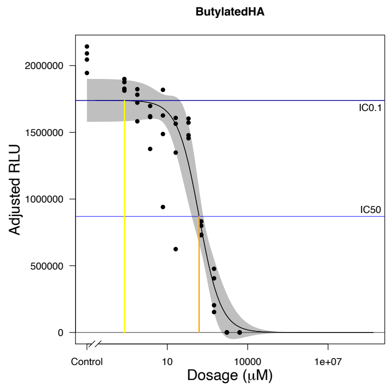
ButylatedHA
2000000
1500000
1000000
500000
0
Adjusted RLU
IC0.1
IC50
Control
10
10000
1e+07
Dosage (μM)

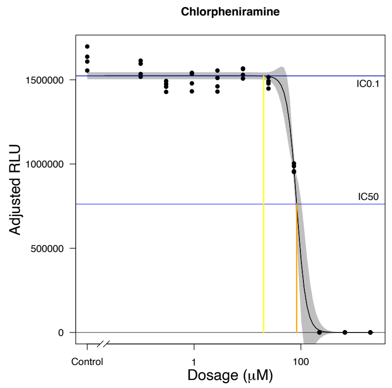
Chlorpheniramine
1500000
1000000
500000
0
Adjusted RLU
IC0.1
IC50
Control
1
100
Dosage (μM)

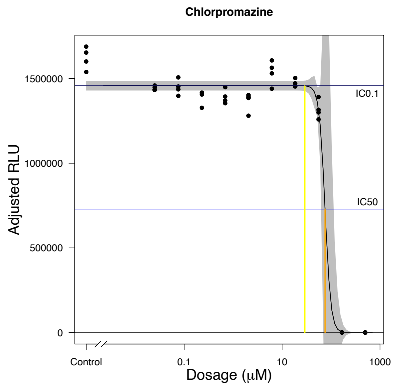
Chlorpromazine
1500000
1000000
500000
0
Adjusted RLU
IC0.1
IC50
Control
0.1
10
1000
Dosage (μM)

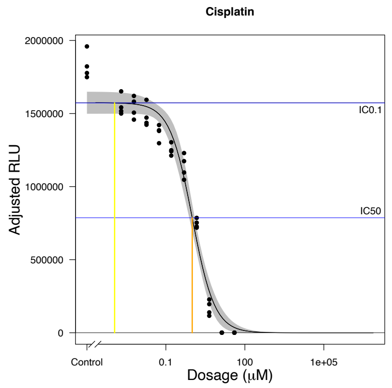
Cisplatin
2000000
1500000
1000000
500000
0
Adjusted RLU
IC0.1
IC50
Control
0.1
100
1e+05
Dosage (μM)

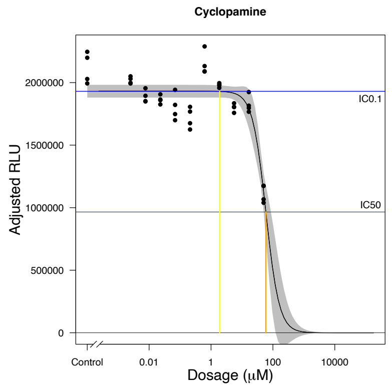
Cyclopamine
2000000
1500000
1000000
500000
0
Adjusted RLU
IC0.1
IC50
Control
0.01
1
100
10000
Dosage (μM)

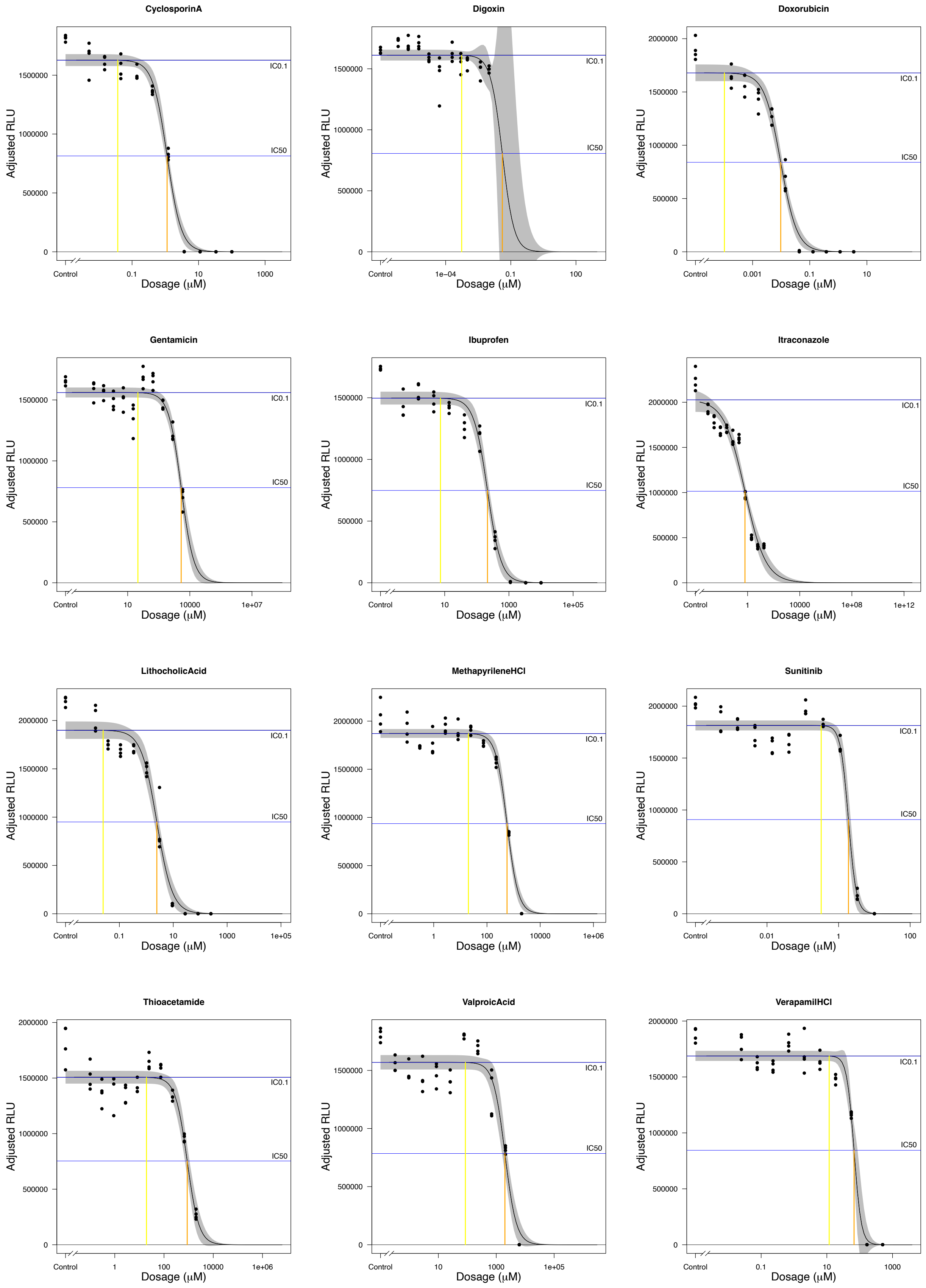


**Fig. S8 Dose response curves of 24 chemicals in the ATP assay using HPS4138 cells**
